# IL-1R signaling drives enteric glia-macrophage interactions in colorectal cancer

**DOI:** 10.1101/2023.06.01.543246

**Authors:** Lies van Baarle, Veronica De Simone, Linda Schneider, Sneha Santhosh, Saeed Abdurahiman, Francesca Biscu, Reiner Schneider, Lisa Zanoletti, Sara Verbandt, Zedong Hu, Michelle Stakenborg, Bo-Jun Ke, Balbina García-Reyes, Jonas Henn, Marieta Toma, Maxime Vanmechelen, Frederik De Smet, Sales Ibiza, Sabine Tejpar, Sven Wehner, Gianluca Matteoli

## Abstract

Enteric glial cells (EGCs) have been implicated in colorectal cancer (CRC) pathogenesis. However, their precise mechanisms of interaction with the CRC immune cell compartment and pro-tumorigenic role remain unclear. This study aimed to investigate the immunomodulatory effects of EGCs on tumor-associated macrophages (TAMs) and their involvement in CRC progression. Using EGC depletion and supplementation models, we assessed the impact of EGCs on the immunomodulation of orthotopic murine CRC. Furthermore, by making use of Bulk RNA-sequencing of CRC EGCs and single-cell sequencing of the tumor microenvironment, we identified the factors involved in the EGC-TAM crosstalk. Findings demonstrate that EGCs acquire a reactive and immunomodulatory phenotype in both murine CRC models and patients, influencing TAM differentiation. Mechanistically, secretion of IL-1 by tumor-infiltrating monocytes and macrophages triggers the phenotypic and functional switch of CRC EGCs via IL-1R. Consequently, tumor EGCs secrete IL-6, promoting the differentiation of monocytes into pro-tumorigenic SPP1^+^ TAMs. Importantly, the reactive tumor EGCs phenotype correlates with worse disease outcomes in preclinical models and CRC patients. Here we uncover a previously unexplored neuroimmune interaction between EGCs and TAMs within the colorectal tumor microenvironment, informing potential therapeutic strategies and enhancing our understanding of CRC progression.

**eTOC Summary:** Our study unveils a novel neuroimmune interaction between enteric glia and TAMs in colon carcinoma. Monocyte/Macrophage-derived IL-1 activates enteric glia, leading to the differentiation of pro-tumorigenic SPP1^+^ TAMs via glial-derived IL-6. Blocking glial IL-1R-signaling reduces colonic tumor lesions, highlighting IL-1R as a potential therapeutic target.

## INTRODUCTION

Identified as the world’s third most common cancer, colorectal cancer (CRC) represents one of the preeminent causes of cancer-associated deaths worldwide. Although innovative technologies have significantly impacted the diagnosis, surgery, and treatment of CRC, patients with advanced disease still have a very poor prognosis. In fact, while the 5-year survival rates of patients with early-stage CRC can reach up to 90%, the survival rate plummets dramatically to as low as 10% for patients diagnosed with advanced metastasis (Kuipers et al., 2015). Hence, a better understanding of the pathogenesis of CRC is crucial to develop new therapeutic strategies along with advanced patient stratification for precision medicine. CRC consists of rapidly evolving neoplasms where acquired mutations in oncogenes and tumor-suppressor genes lead to increasing complexity of the tumor microenvironment (TME), unleashing interaction of the tumor cells with the stroma and the immune system, including fibroblasts, tumor-infiltrating immune cells, and cells of the enteric nervous system (AlMusawi et al., 2021; Albo et al., 2011). This process contributes to the formation of a complex network of cell types within the TME, which is leading to increase tumor fitness.

In recent years, enteric glial cells (EGCs) have also been identified as a new constituent of the colon carcinoma microenvironment (Valès et al., 2019; Yuan et al., 2020). EGCs, once regarded as merely supportive and accessory cells for neurons within the enteric nervous system (Neunlist et al., 2014), have now gained increased attention for their more complex roles in both health and disease (Seguella and Gulbransen, 2021). In homeostasis, EGCs regulate intestinal reflexes and support neurotransmission via communication with enteric neurons. However, accumulating evidence highlights EGCs as crucial mediators of interactions not only among enteric neurons but also intestinal epithelium, enteroendocrine cells, and immune cells (Thomasi and Gulbransen, 2023; Prochera and Rao, 2023; Bohórquez et al., 2014; Seguella and Gulbransen, 2021). Of particular interest is their significant role in modulating immune responses in various intestinal diseases (Ibiza et al., 2016; Progatzky et al., 2021; Grubišić et al., 2020, 2022). Being highly responsive to inflammatory mediators, including ATP, IL-1 cytokines, or LPS, EGCs are rapidly activated during intestinal diseases. Upon activation during intestinal pathologies, EGCs contribute to the shaping of the inflammatory milieu through the secretion of a plethora of cytokines and chemokines (Schneider et al., 2021; Brown et al., 2016; Schneider et al., 2022; Rosenbaum et al., 2016). In this regard, we recently demonstrated the profound influence EGCs exert on macrophage dynamics in the setting of acute intestinal inflammation, promoting the recruitment of monocytes and their differentiation into pro-resolving macrophages (Stakenborg et al., 2022; Schneider et al., 2022).

So far, in the context of CRC, a few studies suggested that EGCs exert a pro-tumorigenic effect during tumor development (Yuan et al., 2020; Valès et al., 2019), however, their influence on the tumor immune microenvironment has not been addressed. In a study by Yuan *et al*., glial cell depletion led to reduced tumor burden in a CRC mouse model (Yuan et al., 2020), indicating a central role of EGCs in CRC development. A xenograft model confirmed this role, and *in vitro* work suggested that EGC activation by IL-1 resulted in a pro-tumorigenic EGC phenotype (Valès et al., 2019), pointing to a direct interaction of EGCs with the TME. However, the mechanisms by which EGCs interact with the different components of the colorectal cancer TME to exert their pro-tumorigenic role remain poorly understood. Especially the molecular and cellular communication pathways involved are so far insufficiently explored and display a substantial lack of *in vivo* evidence.

In this study, we demonstrated, using *in vitro* and *in vivo* models, that upon exposure to the colorectal TME, EGCs undergo a reactive phenotypic switch, leading to the activation of immunomodulatory processes that promote the differentiation of tumor-associated macrophages (TAMs). Intriguingly, tumor-infiltrating monocytes were found to influence the phenotype and function of CRC EGCs through the IL-1 signaling pathway. In turn, EGC-derived IL-6 promoted the differentiation of these monocytes towards SPP1^+^ TAMs. Importantly, this IL-1R/IL-6 axis was found to be essential for the tumor-supportive functions of EGCs.

Together, our findings uncover a novel neuroimmune interaction in the colon cancer microenvironment. This deepens our understanding and may facilitate the development of novel therapeutic approaches to treat this devastating disease.

## MATERIAL AND METHODS

### Animals

WT C57BL/6J (JAX:000664), CCR2^-/-^ (JAX: 004999), PLP1^CreERT2^iDTR (JAX:005975 and JAX:007900), GFAP^Cre^IL-1R1^fl/fl^ (JAX:012886 and JAX:028398), GFAP^Cre^Ai14^fl/fl^ (JAX:012886 and JAX:007908) and Sox10^CreERT2^Ai14^fl/fl^ [Sox10^CreERT2^ (kindly provided by Dr. Vassilis Pachnis (Laranjeira et al., 2011), (Ai14^fl/fl^ JAX:007908)] mice were originally purchased from Jackson Laboratory and bred in our animal facilities. All mice were housed in temperature-controlled specific pathogen–free facilities with ad libitum access to standard chow diet and water under 12-h light–dark cycles at the KU Leuven or University of Bonn. All experimental procedures were approved by the Animal Care and Animal Experiments Ethical Committee of KU Leuven (208/2018 and 213/2018) or by the appropriate authorities of North-Rhine-Westphalia, Germany (81-02.04.2021.A424).

Specific recombination of Sox10^CreERT2^ in enteric glia was confirmed via confocal microscopy by the overlapping tdtomato signal with GFAP immunostaining (Figure S4B). In the case of GFAP^Cre^IL1R1^fl/fl^ mice, following recommendation by The Jackson Laboratory (JAX:012886), we used a strict mating scheme using only Cre^+^ carrying female with Cre^-^ males to overcome any issues of germline Cre-expression and to produce only litters with a GFAP-promotor-driven Cre-expression. Specific GFAP-Cre recombination was confirmed in the reporter mouse line GFAP^Cre^Ai14^fl/fl^ showing in a strong overlap of tdtomato signal with immunolabelled GFAP and SOX10 cells in colonic tissue, confirming Cre activity exclusively in enteric glia (Figure S4E).

### *In vitro* tumor EGCs model

Both orthotopic tumors and healthy colons of C57BL/6J, CCR2^+/+^ or CCR2^-/-^ mice were digested for 30 min in DMEM with 2.5% FBS, 100 µg/mL Penicillin and Streptomycin, 200 U/mL collagenase IV (Gibco, ThermoFisher Scientific) and 125 μg/mL type II dispase (Gibco, ThermoFisher Scientific) to obtain a single-cell suspension. Tumor microenvironment conditioned medium (TME-CM) and healthy colon conditioned medium (H-CM) were generated by culturing 5 x 10^5^ cells/mL in DMEM-complete medium overnight. Next, primary murine embryonic neurosphere-derived EGCs were stimulated with the TME-CM or H-CM for 6, 12 or 24 hours. For IL-1R blocking experiments, primary embryonic neurosphere-derived EGCs were incubated for 24h with TME-CM together with 5 µg/mL isotype IgG (BioXCell) or 5 µg/mL anti-IL-1R (BioXCell).

### Orthotopic CRC model

Orthotopic colonic sub-mucosal implantation of CRC cells was performed as previously described (Zigmond et al., 2011). Briefly, MC38 cells (Corbett et al., 1975) were intracolonic (i.c.) injected via endoscopy as a single-cell suspension containing between 75 000 – 750 000 MC38 cells/ 100 µL PBS depending on the susceptibility of the mouse strain. For the EGCs supplementation model, primary embryonic neurosphere-derived EGCs were isolated with 0.05% Trypsin-EDTA (Gibco, ThermoFisher Scientific) and treated with HBSS (Gibo, ThermoFisher Scientific) supplemented with 100 µg/mL DNAse I (Roche) and 5 mM MgCl_2_ (Sigma-Aldrich) for 30 min at RT. Subsequently, EGCs were first washed with HBSS with 5 mM MgCl_2_ and then with PBS. Finally, EGCs were resuspended in PBS together with MC38 cells in a ratio of 1:1 and orthotopically co-injected in C57BL/6J WT mice. Two weeks prior to the start of each experiment, PLP1^CreERT2^iDTR mice were injected intraperitoneally (i.p.) 2 times every other day with 100 mg/kg Tamoxifen (Sigma-Aldrich) dissolved in 100 µL MIGLYOL®812 (Sigma-Aldrich). For EGCs *in vivo* depletion experiments, PLP1^CreERT2^iDTR mice were injected i.c. with 2 mg/kg Diphtheria toxin (DT) (Merck, Sigma) dissolved in 100 µL of saline, three and five days prior to the start of the tumor implantation. Tumor volume was determined by caliper measurements and calculated based on the height (h), length (l) and width (w) of the tumor, according to the formula: (π/6)*h*l*w.

### AOM-DSS model

Female GFAP^Cre^IL-1R1^fl/fl^ and GFAP^Wt^IL-1R1^fl/fl^ littermate mice (10-14 weeks of age) were injected i.p. with azoxymethane (AOM; 10 mg/kg; Sigma-Aldrich) a week prior starting 3 cycles of DSS colitis using 1.5% DSS in drinking water (MP Biomedicals) for 5 days followed by 16 days of recovery with normal drinking water (Parang et al., 2016). On day 70 colonic tumor development was determined.

## RESULTS

### EGCs shape the CRC immune compartment

Recent studies identified EGCs as an important component of the colon TME. However, their contribution to CRC pathogenesis and their possible interaction with the tumor immune compartment remains largely unexplored. Hence, to address the pro-tumorigenic and immunomodulatory role of EGCs, MC38 murine colorectal cancer cells were orthotopically injected into the colonic submucosa (Figure 1A and Figures S1A-S1B) of PLP1^CreERT2^iDTR mice allowing temporal and local depletion of enteric glia during colon tumor development. Following tamoxifen exposure, diphtheria toxin (DT) was delivered by colonoscopy-guided injections into the colonic wall on days –5 and –3 before submucosal MC38 injection (Figure 1B). Mucosal injection of DT resulted in local EGC depletion, as indicated by decreased GFAP protein level at the injection site while normal GFAP protein levels were observed 1.5 cm adjacent to the DT injection site, with overall discernible impact on the total colon length (Figures S1C-S1E). However, PLP1 reporter-guided EGCs depletion only leads to a temporary depletion, as EGCs have been reported to repopulate the gut within 2 weeks following the initial depletion (Baghdadi et al., 2021). Taking advantage of the orthotopic CRC model, which confers precise control of tumor location and growth rate (Zigmond et al., 2011), the endpoint of our orthotopic depletion experiment was chosen 7 days post-tumor induction (a total of 12 days after the first DT injection). Seven days after colonic MC38 cells injection, a significant reduction of tumor size was observed in DT pre-treated mice compared to the vehicle group (Figure 1C). Interestingly, during this early phase of tumor growth (7 days), the depletion of EGCs resulted also in a decreased number of TAMs (Figure 1D and S1F). Furthermore, a decrease in the numbers of monocytes and eosinophils was observed in tumors after enteric glia depletion, whereas no differences were observed for neutrophils, T and B cells (Figures 1D and S1G).

**Figure 1.**
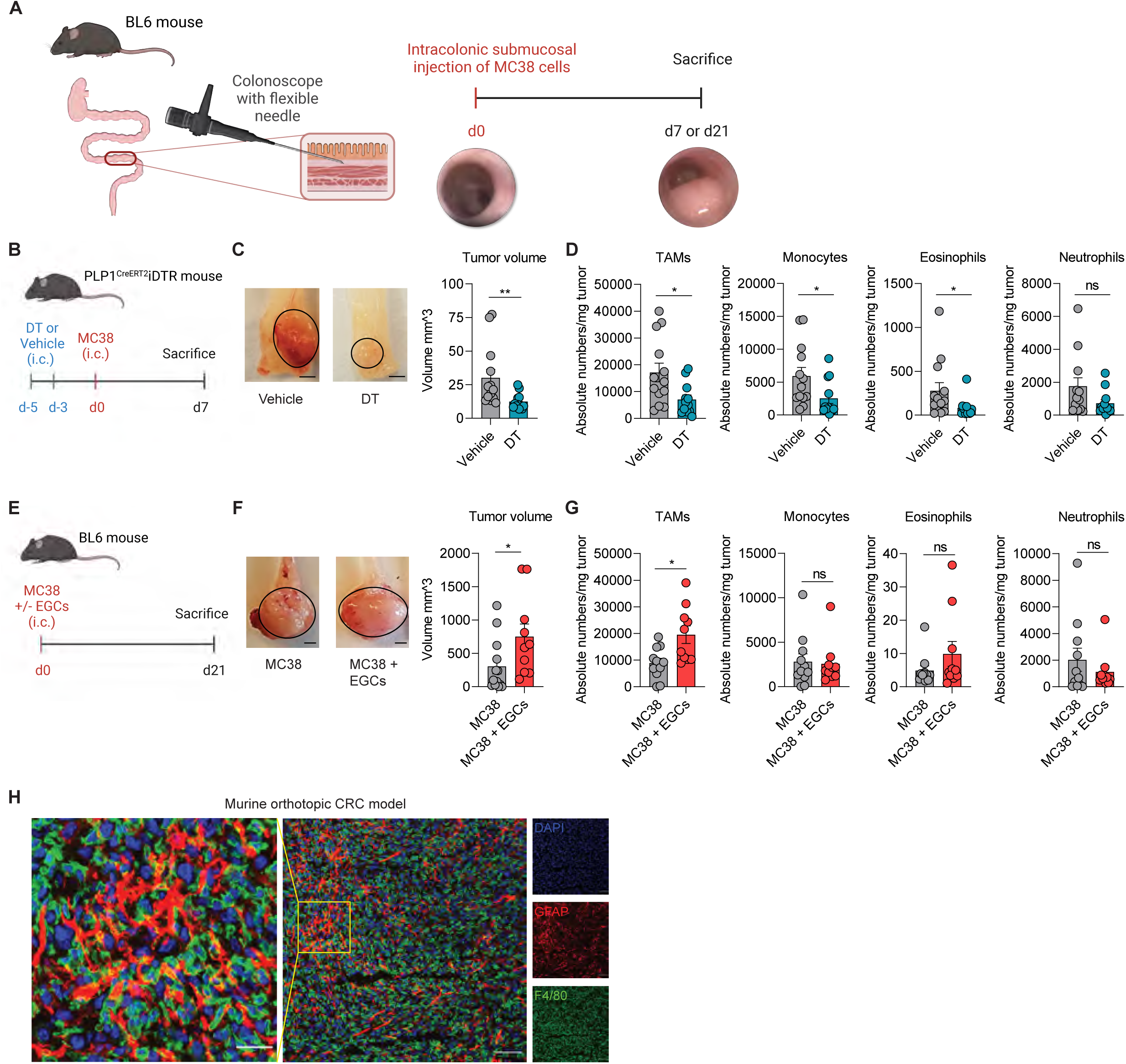
EGCs shape the CRC immune compartment. (A) BL6 mice were injected endoscopically in the colonic submucosa at day(d)0 with MC38 cells. Tumors were assessed at d7 or d21. Schematic representation of the murine orthotopic CRC model. (B-D) PLP1^CreERT2^iDTR mice were intracolonic (i.c.). at d-5 and d-3 with 40 ng Diphtheria toxin (DT) or saline (Vehicle). At d0, MC38 cells were i.c. injected in both groups. Tumor growth and myeloid immune infiltration were assessed at d7. Schematic representation of EGCs depletion mouse model (B) with representative pictures (scale bar 2 mm) and quantitative comparison of tumor volume (C). Data show absolute tumor-infiltrating myeloid immune cell numbers per mg tumor tissue (D) (*n = 13* Vehicle, *n = 12* DT). (E-G) BL6 mice were intracolonic injected at d0 with MC38 cells with or without embryonic neurosphere-derived EGCs (1:1 ratio). Tumor growth and myeloid immune infiltration were assessed at d21. Schematic representation of EGCs co-injection mouse model (E) with representative pictures (scale bar 2 mm) and quantitative comparison of tumor volume (F). Data show absolute tumor-infiltrating myeloid immune cell numbers per mg tumor tissue (G) (*n = 11* MC38, *n = 10* MC38 + EGCs). (H) Representative image showing GFAP (red), F4/80 (green) and DAPI (blue) in orthotopic murine tumor sections, (scale bar 70 µm and 25 µm). Data show mean ± SEM. Statistical analysis: unpaired Mann-Whitney test (C-D,F-G) *p < 0.05, **p < 0.005, ns not significant. See also Figure S1.

To further validate the effect of EGCs on the CRC immune compartment, we established a co-injection model of MC38 cells together with primary EGCs (Figure 1E). Co-injection of MC38 and EGCs resulted in increased tumor growth together with higher numbers of TAMs, as well as CD4^+^ T cell, CD8^+^ T cell, and T_reg_ cells compared to mice orthotopically injected with MC38 cells alone (Figures 1F-1G and S1H). No differences were observed in the numbers of monocytes, eosinophils, neutrophils, and B cells (Figures 1G and S1H).

Spatial tissue mapping via confocal microscopy conformed close proximity between EGCs (GFAP^+^) and TAMs (F4/80^+^) within orthotopic colonic tumors injected in WT mice (Figure 1H), further suggesting the existence of a glial-immune interplay within the TME.

Overall, these findings suggest that EGCs participate in shaping the CRC immune microenvironment, by expanding the TAM population.

### EGCs display an activated and immunomodulatory phenotype in CRC

To examine the molecular mechanisms by which EGCs influence the immune CRC compartment with particular regard to the TAMs, we first investigated their transcriptional adaptations upon CRC onset. To this end, we established an *in vitro* tumor EGC model, able to mimic the response of enteric glia to the factors secreted by the colonic tumor microenvironment (TME). To this end, primary embryonic-derived EGCs were treated with conditioned medium (CM) of digested murine MC38 orthotopic tumors, from now onwards, defined as TME-CM EGCs (Figure 2A). In this model, EGCs stimulated with supernatants derived from healthy colonic tissue (H-CM) and naïve unstimulated EGCs were used as controls. At 6 hours, 12 hours, and 24 hours post-stimulation bulk RNA sequencing (RNA-seq) was analyzed uncovering strong transcriptional differences among the various EGC groups. Principal component analysis (PCA) demonstrated a significant similarity between the H-CM and unstimulated EGC samples (Figure 2B). In contrast, TME-CM EGCs exhibited a distinct separation from both H-CM and unstimulated EGCs, suggesting a noticeable difference in their transcriptional programs. Next, using weighted gene correlation network analysis (WGCNA) we identified 12 gene co-expression modules. TME-CM EGCs showed specific correlation to modules 7 and 8 and an inverse correlation to module 4 (Figure 2C; Table S1). Here, module 7 showed a functional association with glial reactivity, indicated by the upregulation of genes such as *Lcn2* and *Timp1*, which are typical markers for pan-reactive astrocytes (Clarke et al., 2018) (Figure 2D). Module 8 consisted of genes, including *Ccl2* and *Il6,* that are associated with immunomodulatory functions of EGCs, mimicking a state of enteric gliosis (Schneider et al., 2022). Conversely, the genes of module 4, such as *Ntsr1* and *Sparlc1,* were associated with the homeostatic functions of EGCs (Drokhlyansky et al., 2020). In line, gene set enrichment analysis of the 24h TME-CM EGCs signature revealed impairment for functions previously ascribed to healthy EGCs, including GO terms like “*Positive regulation of stem cell differentiation”, “Regulation of glial cell differentiation and gliogenesis”, “Neuron projection guidance”,* and *“Positive regulation of neurogenesis”* (Baghdadi et al., 2021; Seguella and Gulbransen, 2021) (Figure 2E; Table S2). Notably, in line with the previous *in vitro* findings of Valès *et al*., TME-CM EGCs were also enriched for the GO terms such as “*Positive regulation of prostaglandin biosynthetic process”,* and “*Interleukin 1 receptor activity”*, supporting the possible paracrine effect of IL-1/PGE_2_ signaling (Valès et al., 2019). Lastly, gene set enrichment analysis predicted a direct interaction of CRC EGCs with TAMs, reflected by functional enrichment for the GO terms “*Macrophage differentiation”* and *“Positive regulation of macrophage activation* and *migration”* (Figure 2E). Taken together, upon exposure to the CRC TME, EGCs undergo a phenotypic switch associated with the activation of immunomodulatory programs related to macrophage interplay.

**Figure 2.**
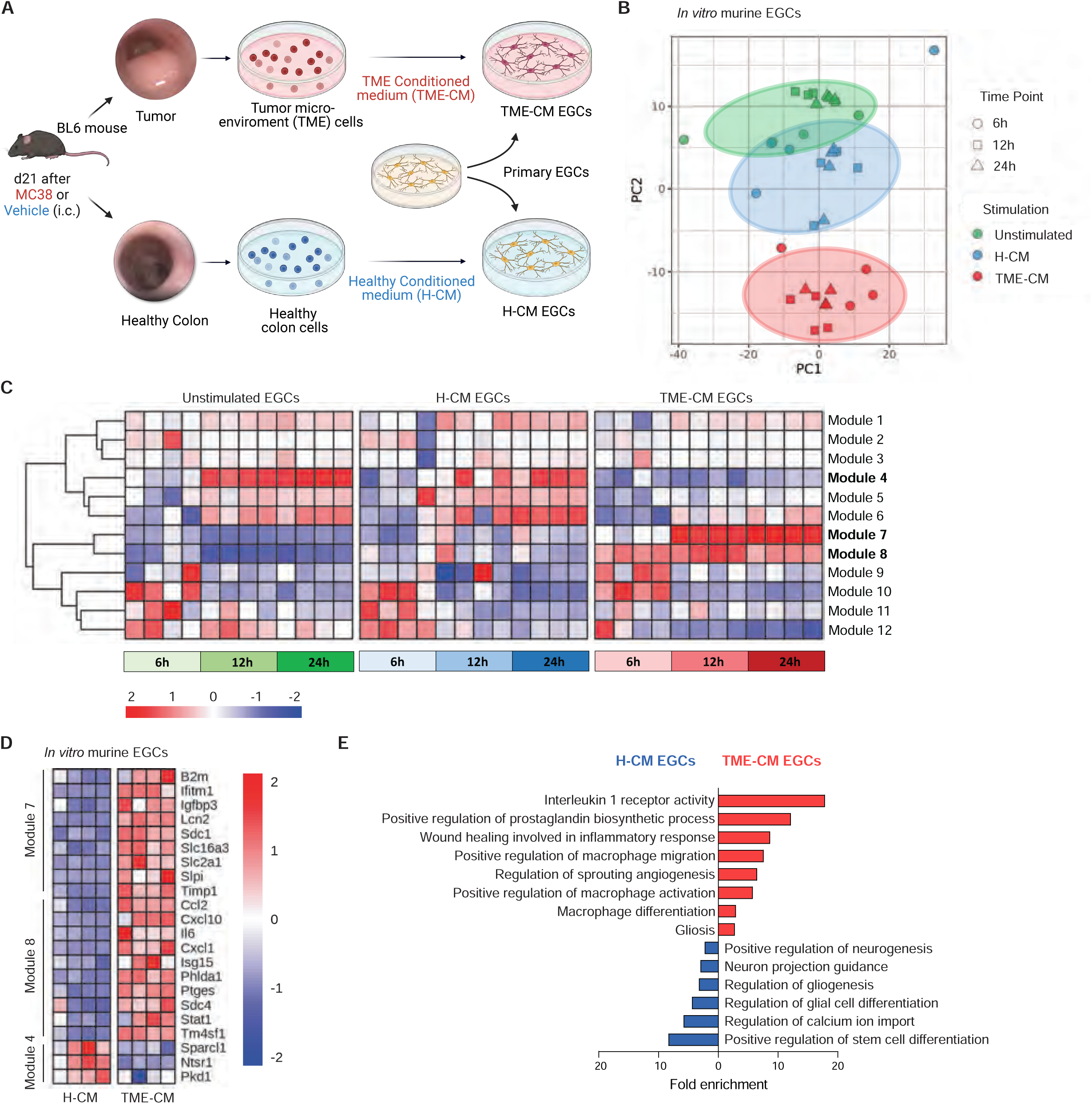
EGCs display an activated and immunomodulatory phenotype in CRC. Transcriptome analysis of *in vitro* primary embryonic neurosphere-derived EGCs alone or stimulated with healthy conditioned medium (H-CM) or tumor microenvironment conditioned medium (TME-CM) at different time points (6h, 12h, and 24h, *n = 4*). (A) Schematic representation of the *in vitro* tumor EGCs model. (B) Principal component analysis (PCA) plot of EGCs gene signature identified by 3’mRNA bulk RNA-seq. Each dot represents an individual sample. (C) Heatmap showing the transcriptional modules identified by weighted gene correlation network analysis (WGCNA). (D) Heatmap of differentially expressed genes in modules 4, 7, and 8 of *in vitro* murine EGCs stimulated for 24h with H– or TME-CM. (E) Gene set enrichment analysis for the differentially up– and down-regulated genes in TME-CM versus H-CM stimulated EGCs after 24h (*n = 4*). See also Table S1 and S2.

### Tumor EGC-derived IL-6 favors SPP1^+^ TAM differentiation

Considering that tissue location and transcriptomic data suggest direct communication between EGCs and TAMs in the CRC microenvironment, we aim to decipher the molecular mechanisms underpinning their interaction. Firstly, single-cell transcriptomics was used to characterize the immune landscape of colorectal MC38 orthotopic tumors (Figure S2A). Interestingly, among the identified immune populations, monocytes and macrophages accounted for 60% of the tumor-infiltrating immune cells. Unsupervised clustering of the myeloid cells (*Lyz2, Cd68, H2-Ab1, Mrc1, C1qa, Ly6c2, Ccr2,* and *Fn1*) revealed 1 monocyte and 4 distinct macrophage sub-populations (Figures 3A and 3B). The macrophage cluster that was most abundant was characterized by marker genes both for monocytes (Ccr2 and Ly6c2) and for differentiated macrophages (H2-Ab1 and Nlrp3). This suggests a possible transitional state, which we have termed ‘Intermediate Macrophages’ (Figure 3B). Additionally, we identified two clusters of TAMs, ‘SPP1^+^ TAMs’ co-expressing of *Spp1* and *Arg1,* together with genes involved in angiogenesis (*Vegfa*) and extracellular matrix remodeling (*Spp1* and *Tnf*) (Figures 3B and S2B) together with ‘C1Q^+^ TAMs’ characterized by genes involved in phagocytosis (*Nr1h3*), antigen presentation (*H2-Ab1*) and the complement cascade (*C1qa*) (Figures 3B and S2B). A second C1Q^+^ TAM cluster, characterized by high expression of cell cycle genes, including *Mki67* and *Top2a,* was classified as ‘Cycling C1Q^+^ TAMs’ (Figure 3B).

**Figure 3.**
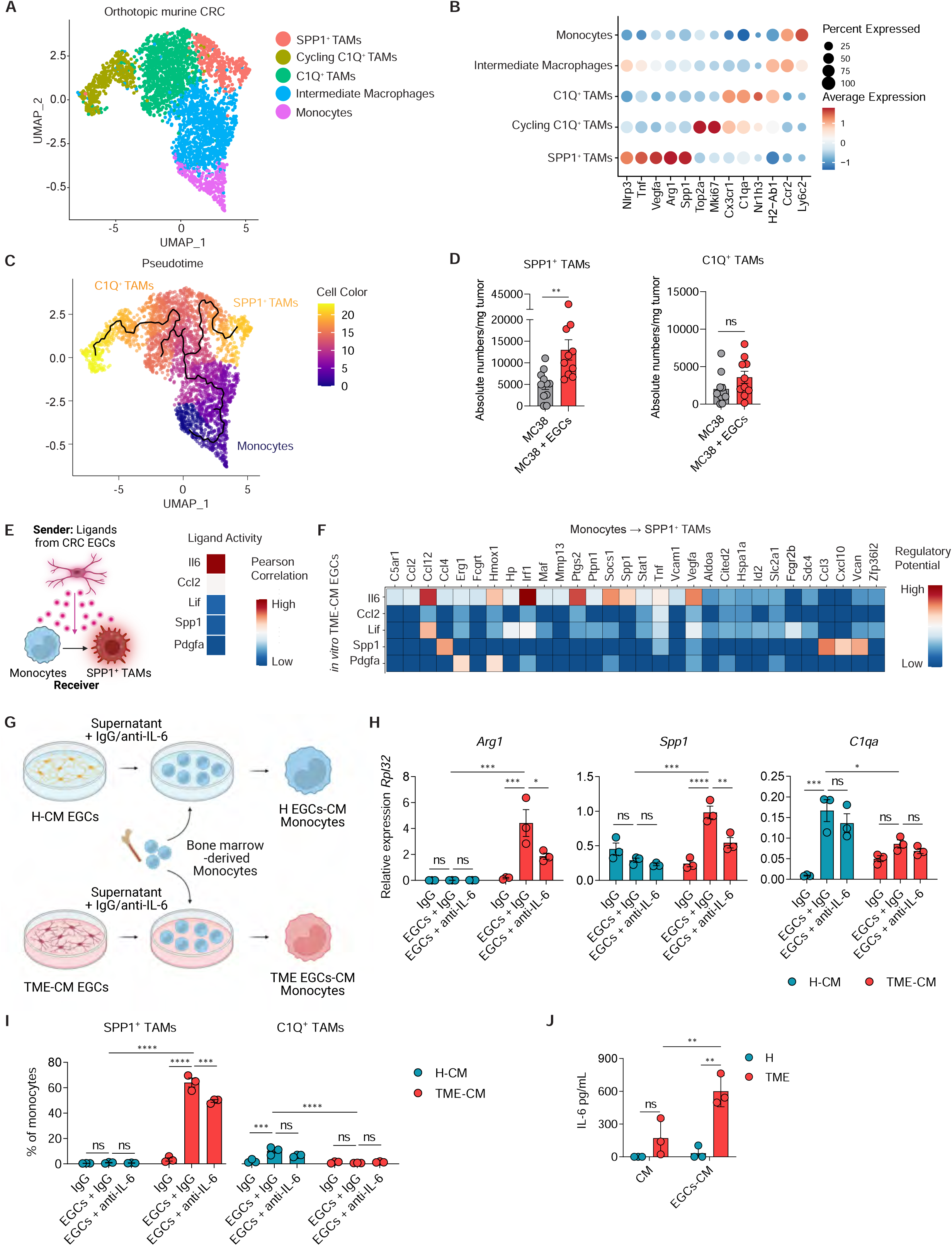
Tumor EGC-derived IL-6 favors SPP1^+^ TAM differentiation. (A-C) scRNA-seq analysis of monocytes and macrophages from the scRNA-seq data of tumors from BL6 mice bearing orthotopic MC38 tumors, 21 days(d) after tumor induction *(n = 3)*. UMAP of tumor-infiltrating monocyte and macrophage subclusters (A) and dot plot of differentially expressed marker genes used for their annotation (B). Differentiation trajectory of murine CRC-infiltrating monocyte and macrophage subsets inferred by Monocle (C). (D) BL6 mice were intracolonic injected with MC38 cells and with or without embryonic neurosphere-derived EGCs (1:1 ratio). SPP1^+^ TAMs and C1Q^+^ TAMs infiltration was assessed on d21. Data represents absolute numbers per mg tumor tissue (*n = 11* MC38, *n = 10* MC38 + EGCs). (E-F) Nichenet analysis was performed considering the ligands expressed by 24h TME-CM EGCs, data extracted from *in vitro* CRC EGCs bulk RNAseq dataset (See Figure 2) and considering the differentially expressed genes between monocytes and SPP1^+^ TAMs as target genes, data extracted from CRC orthotopic murine CRC dataset (See Figure 3A). Top ligands from *in vitro* tumor microenvironment conditioned medium (TME-CM) EGCs and their Pearson correlation, predicted to be inducing monocytes to SPP1^+^ TAM differentiation according to NicheNet (E). Heatmap of ligand-target pairs showing regulatory potential scores between top positively correlated prioritized ligands of *in vitro* TME-CM EGCs and their target genes among the differentially expressed genes between monocytes and SPP1^+^ TAMs (F). (G-I) *In vitro* murine bone marrow-derived monocytes cultured for 48h with supernatant of healthy (H)-CM, TME-CM, H EGCs-CM, and TME EGCs-CM in the presence of IgG or anti-IL-6 (both 5 µg/mL) *(n = 3)*. Experimental design (G). Relative mRNA levels for *Arg1, Spp1,* and *C1qa* normalized to the housekeeping gene *Rpl32* in monocytes after stimuli (H). Percentages of SPP1^+^ and C1Q^+^ TAMs in monocyte cultures after stimuli (I). (J) IL-6 concentration in the H-CM, TME-CM, H EGCs-CM, and TME EGCs-CM *(n = 3)*. All data are represented as mean ± SEM. Statistical analysis: unpaired t-test (D), two-way ANOVA test with correction for multiple comparisons (H-J). * p <0.05, ** p <0.005, *** p <0.0005, **** p <0.00005, ns not significant. See also Figure S2.

Overall, our findings are in line with the study of Zhang *et al*., which reported very similar dichotomous functional phenotypes of TAMs in CRC patients (Zhang et al., 2020). Additionally, Zhang and colleagues predicted a dichotomic differentiation trajectory of monocytes towards SPP1^+^ TAMs or C1QC^+^ TAMs in CRC patients. In line, we identified a strong directional flow from tumor-infiltrating monocytes towards intermediate macrophages, which in turn further branched into two opposite paths, ending either in SPP1^+^ TAMs or C1Q^+^ TAMs (Figure 3C).

SPP1^+^ macrophages represent a significant cell population within the CRC immune cell compartment and are considered potential prognostic markers in CRC. However, the current knowledge regarding the microenvironmental cues, promoting the differentiation of tumor-infiltrating monocytes towards SPP1^+^ TAMs or C1Q^+^ TAMs, remains limited. Thus, we explored the possible role of EGCs in promoting monocyte to SPP1^+^ TAM or C1Q^+^ TAM differentiation using our EGC co-injection CRC model (Figures 3D and S2C). Strikingly, supplementation of EGCs within the CRC TME resulted in more than a twofold increase in SPP1^+^ TAMs, while no significant difference was found for C1Q^+^ TAMs. Next, to identify key EGC-derived mediators accountable for SPP1^+^ TAM differentiation in CRC, we applied NicheNet, a computational tool designed to infer relationships between signaling molecules and their target gene expression (Browaeys et al., 2020). By using the genes differentially expressed between SPP1^+^ TAMs and monocytes [data extracted from our single-cell RNA-seq (scRNAseq) murine orthotopic CRC dataset] as target genes, we prioritized candidate ligands derived from TME-CM EGCs (data extracted from bulk RNAseq *in vitro* CRC EGCs dataset) as potential drivers of this differentiation process (Figure 3E). Here, TME-CM EGC-derived IL-6 was identified as the top predicted candidate factor involved in driving the SPP1^+^ TAM phenotype (Figures 3E and 3F).

In line with our prediction, IL-6 neutralization in TME-CM EGC supernatant (Figure 3G) attenuated the differentiation of monocytes into SPP1^+^ TAMs, further reflected by reduced *Spp1* and *Arg1* expression (Figures 3H and 3I). Of note, TME-CM EGCs did not promote C1Q expression in monocytes, suggesting that CRC EGCs specifically favor SPP1^+^ over C1Q^+^ TAM differentiation (Figures 3H and 3I). Consistent with this, quantification of IL-6 revealed elevated levels in the supernatant of TME-CM EGCs compared to H-CM EGCs (Figure 3J).

Altogether, our data highlight an important and previously overlooked interaction between EGCs and TAMs in the CRC TME, where EGC-derived IL-6 might be a key regulator for driving SPP1^+^ TAM differentiation.

### Monocyte-derived IL-1 promotes the CRC EGC phenotype

Given the evidence that EGCs modulate their functions based on microenvironmental cues (Progatzky et al., 2021; Stakenborg et al., 2022; Schneider et al., 2021), we aimed at defining the factors driving the EGC phenotypic switch in CRC. Considering the heavy infiltration of the colon TME by immune cells, which recent studies have pinpointed as sources of EGC-activating factors (Progatzky et al., 2021), we investigated the possibility of immune cells driving CRC EGC transition, potentially creating a reinforcing neuroimmune feedback loop. To investigate the cellular circuits coordinating this interaction we used NicheNet (Browaeys et al., 2020) linking ligands derived from immune cells within the TME (from the scRNAseq murine orthotopic CRC dataset) to target genes differentially expressed between H-CM EGCs and TME-CM EGCs (data extracted from Bulk RNAseq *in vitro* CRC EGCs dataset) (Figure 4A). In this analysis, IL-1β and IL-1α emerged as the top-ranked ligands driving the transcriptional transition from H-CM EGCs to TME-CM EGCs (Figure 4A and 4B). Notably, we could confirm elevated levels of IL-1 in the TME samples compared to healthy colon cells, both at as RNA and protein (Figure 4C and S3A). Consistent with our prediction, treating primary EGCs with IL-1β markedly activated the CRC EGC phenotype, with induction of Lcn2 and Timp1, and immunomodulatory factors Ccl2 and Il6, both at RNA and protein levels (Figures 4D-4E, Figures S3B-S3C; Table S2). Conversely, inhibiting IL-1R signaling in EGCs during stimulation with the TME-CM, completely abrogated the induction of the CRC EGC key markers (Figure 4F and 4G). Taken together, these findings underscore the pivotal role of IL-1 in the reprogramming of EGCs upon exposure to the colonic TME.

**Figure 4.**
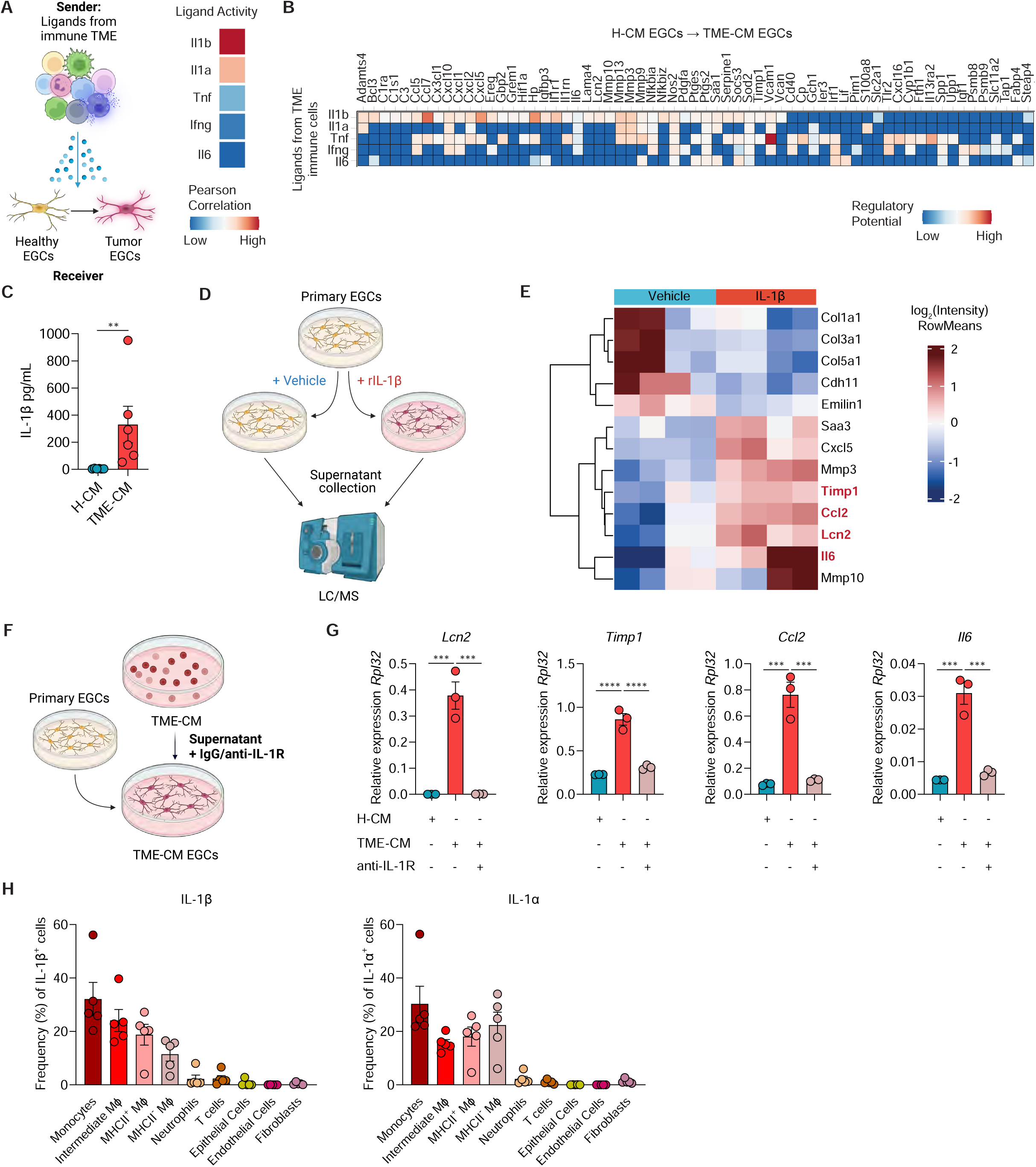
IL-1R activation drives the CRC EGC phenotype. (A-B) Top ligands from orthotopic CRC tumor-infiltrating immune cells predicted by NicheNet to be inducing CRC EGC signature. Nichenet analysis was performed considering the ligands expressed by murine orthotopic CRC tumor-infiltrating immune cells, data extracted from CRC orthotopic murine CRC dataset (See Figure S2A) and considering the differentially expressed genes between 24h Healthy conditioned medium (H-CM) EGCs and tumor microenvironment (TME)-CM EGCs as target genes, data extracted from *in vitro* CRC EGCs bulk RNAseq dataset (See Figure 2). Schematic representation of TME-derived ligand-EGC interplay (left) and top 5 predicted ligands with their Pearson correlation (right) (A). Heatmap of ligand-target pairs showing regulatory potential scores between top ligands and target genes among the differentially expressed genes between *in vitro* H-CM EGCs and TME-CM EGCs (B). (C) Protein level of IL-1β in H-CM and TME-CM *(n = 6)*. (D-E) Primary adult neurosphere-derived EGCs were isolated from BL6 mice and treated with or without rIL-1β (10 ng/mL) for 24h. Protein concentration in the culture supernatants was determined by liquid chromatography/mass spectrometry (*n = 4*). Schematic experimental representation (D) and heatmap of differentially expressed proteins between Vehicle and IL-1β-treated EGCs (E). (F-G) Primary embryonic neurosphere-derived EGCs were stimulated for 24h with H-CM or TME-CM in the presence or absence of IgG or anti-IL-1R (5 µg/mL each) *(n = 3)*. Schematic representation of experimental setup (F). Relative mRNA levels for *Lcn2, Timp1, Ccl2,* and *Il6* normalized to the housekeeping gene *Rpl32* (G). (H) BL6 mice were intracolonic injected at day(d)0 with MC38 cells, and both stromal and immune cells were assessed for IL-1β and IL-1α expression at d21. Data are presented as the frequency of total live IL-1β^+^ or IL-1α^+^ cells *(n = 5).* All data are represented as mean ± SEM (C, G-H). Statistical analysis: unpaired Mann-Whitney test (C), unpaired t-test (G). ** p <0.005, *** p<0.0005, **** p <0.00005 See also Figure S3 and Table S2.

Next, to identify the cellular source of IL-1 within the TME, we quantified IL-1 expression across epithelial, stromal, and immune cells (Figures 4H and S3D-S3F). While Valés *et al*. concluded that *in vitro* IL-1 is released by the tumor epithelial cells, we herein demonstrate that *in vivo* in the TME IL-1 secretion is restricted to myeloid cells, while no expression could be found in epithelial, nor in stromal cells (Figure S3F). Further analyses identified tumor monocytes as the principal producers of both IL-1β and IL-1α at the RNA and protein levels (Figures 4H and S3G).

To define the possible effect of tumor monocyte-derived IL-1 on the transcriptional reprogramming of EGCs, we isolated tumor– and bone marrow (BM)-derived monocytes from mice bearing orthotopic colon tumors and exposed primary enteric glia to their supernatant with or without IL-1R blockade (Figure 5A). Remarkably, the supernatant of tumor monocytes was able to induce a higher expression of CRC EGC marker genes (*Lcn2*, *Timp1*, *Ccl2*, and *Il6*) compared to control BM-derived monocytes in an IL-1R-dependent manner (Figure 5B).

**Figure 5.**
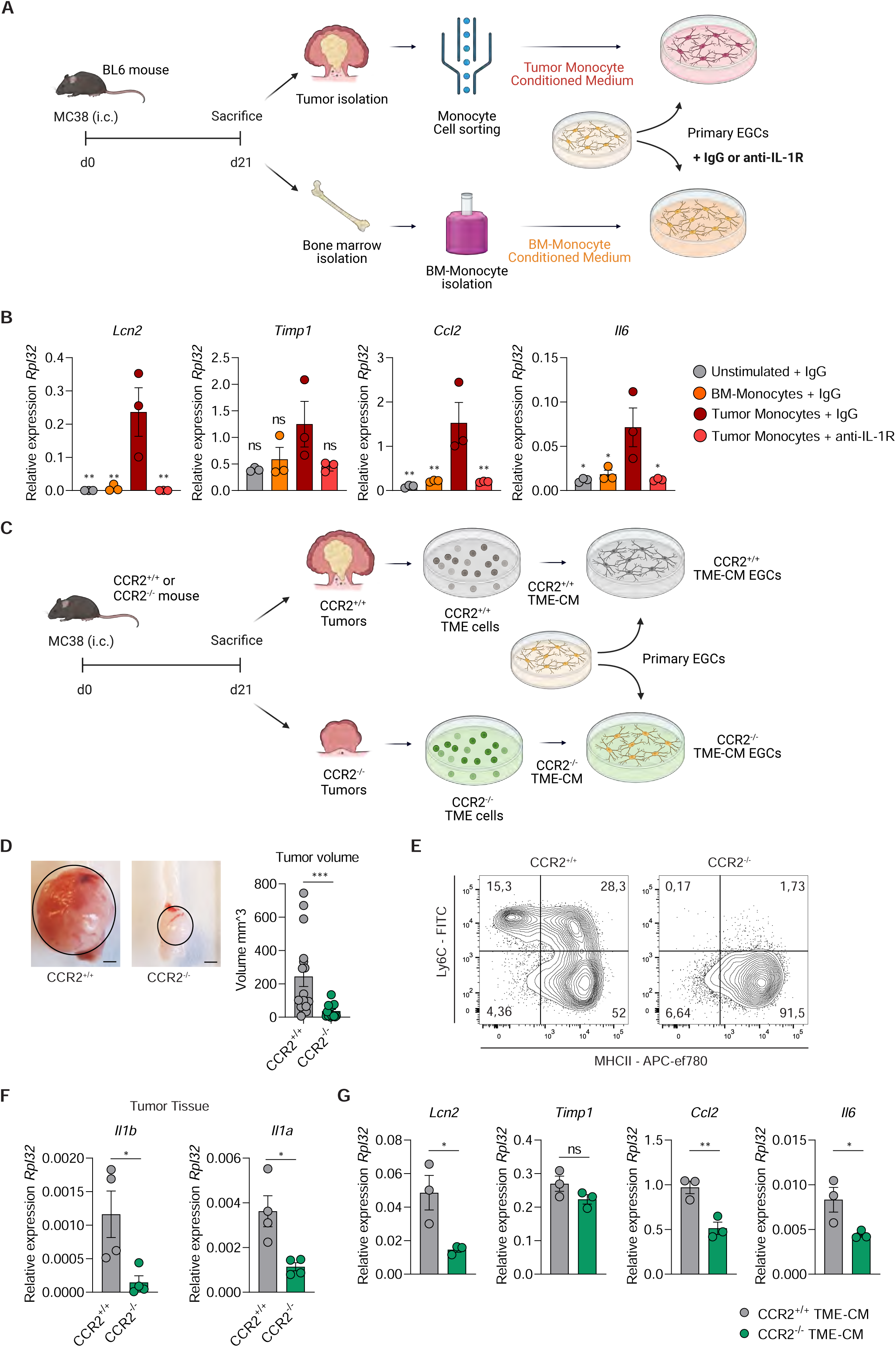
Monocyte-derived IL-1 promotes the CRC EGC phenotype. (A-B) Primary embryonic neurosphere-derived EGCs were stimulated for 24h with IgG or anti-IL-1R (5 µg/mL each) with or without the supernatant of sorted tumor monocytes or bone marrow (BM)-derived monocytes from BL6 mice bearing orthotopic CRC tumors. Schematic representation of experimental setup (A). Relative mRNA levels of *Lcn2, Timp1, Ccl2,* and *Il6,* normalized to the housekeeping gene *Rpl32,* in primary embryonic neurosphere-derived EGCs were compared between EGCs stimulated with tumor monocyte supernatant and all other conditions *(n = 3)* (B). (C-G) CCR2^+/+^ and CCR2^-/-^ mice were intracolonic injected at day(d)0 with MC38 cells, tumor tissue was collected at d21. Then, *in vitro* embryonic neurosphere-derived EGCs were cultured for 24h with the tumor microenvironment-conditioned medium (TME-CM) of CCR2^+/+^ and CCR2^-/-^ tumors. Schematic representation of experimental setup (C). Representative pictures (left, scale bar 2 mm) and quantitative comparison of tumor volume (right) (*n = 16* CCR2^+/+^, *n = 17* CCR2^-/-^) (D). Representative contour plots of tumor-infiltrating monocytes and macrophages gated on live-CD45^+^-CD11b^+^-Ly6G^−^– CD64^+^ cells (E). Relative mRNA levels of *Il1b* and *Il1a* normalized to the housekeeping gene *Rpl32* in CCR2^+/+^ and CCR2^-/-^ CRC tumors *(n = 3)* (F). Relative mRNA levels of *Lcn2, Timp1, Ccl2,* and *Il6* in EGCs stimulated with TME-CM of CCR2^+/+^ and CCR2^-/-^ mice, normalized to the housekeeping gene *Rpl32 (n = 3)* (G). Data represented as mean ± SEM (B, D, F-G). Statistical analysis: One-way ANOVA test with correction for multiple comparisons, compared to tumor monocyte supernatant + IgG condition (B), Mann Whitney test (D) or unpaired t-test (F-G). *p < 0.05, ** p <0.005, *** p <0.0005, ns not significant.

To further verify the monocyte origin of IL-1 signaling, we examined the effects on primary EGCs of orthotopic TME-CM, sourced from tumors induced in both monocyte-deficient [C-C chemokine receptor type 2 deficient (CCR2^-/-^)] and monocyte-competent mice (CCR2^+/+^) (Figure 5C). Consistent with previous findings (Afik et al., 2016), the volume of orthotopic tumors grown in the colonic mucosa of CCR2^-/-^ mice was significantly reduced (Figures 5D) as consequence of the reduce number of recruited monocyte and monocyte-derived macrophage in CCR2^-/-^ tumors as confirmed by flow cytometry (Figure 5E). Consistently, *Il1b* and *Il1a* expressions were significantly decreased in the tumor tissue of CCR2-deficient mice compared to WT mice (Figure 5F), further corroborating that monocytes and monocyte-derived macrophages are the major source of IL-1 ligands in the colon TME. As expected, TME-CM isolated from the CCR2^-/-^ mice failed to induce CRC EGC reprogramming as reflected by the reduced expression of *Lcn2, Ccl2,* and *Il6* when compared with EGCs treated with CCR2^+/+^ TME-CM (Figure 5G). A similar trend, although not significant, was observed for *Timp1*. Overall, our findings strongly support the concept that IL-1, derived from tumor-infiltrating monocytes– and monocyte-derived macrophages, provides remodeling of the neighboring enteric glia into activated and immunomodulatory CRC EGCs.

### IL-1R signaling in EGCs promotes SPP1^+^ TAM differentiation and tumor progression

Next, we assessed whether IL-1R blocking in CRC EGCs might directly affect TAM differentiation. For this purpose, EGC cultures were exposed to TME-CM or H-CM in the presence of an IL-1R blocking antibody and subsequently their supernatant was used to treat naive monocytes (Figure 6A). The blockade of IL-1R reduced the differentiation of monocytes into SPP1^+^ TAMs in the context of TME-CM EGCs, as evidenced by the reduced expression of SPP1 and ARG1 (Figures 6B and S4A). As a result, IL-6 levels were markedly reduced in the supernatant of TME-CM-exposed EGCs following IL-1R inhibition (Figure 6C). These findings further corroborate our hypothesis regarding the critical role of the IL-1R/IL-6 axis in CRC EGCs in directing monocyte differentiation towards the SPP1^+^ TAM phenotype.

**Figure 6.**
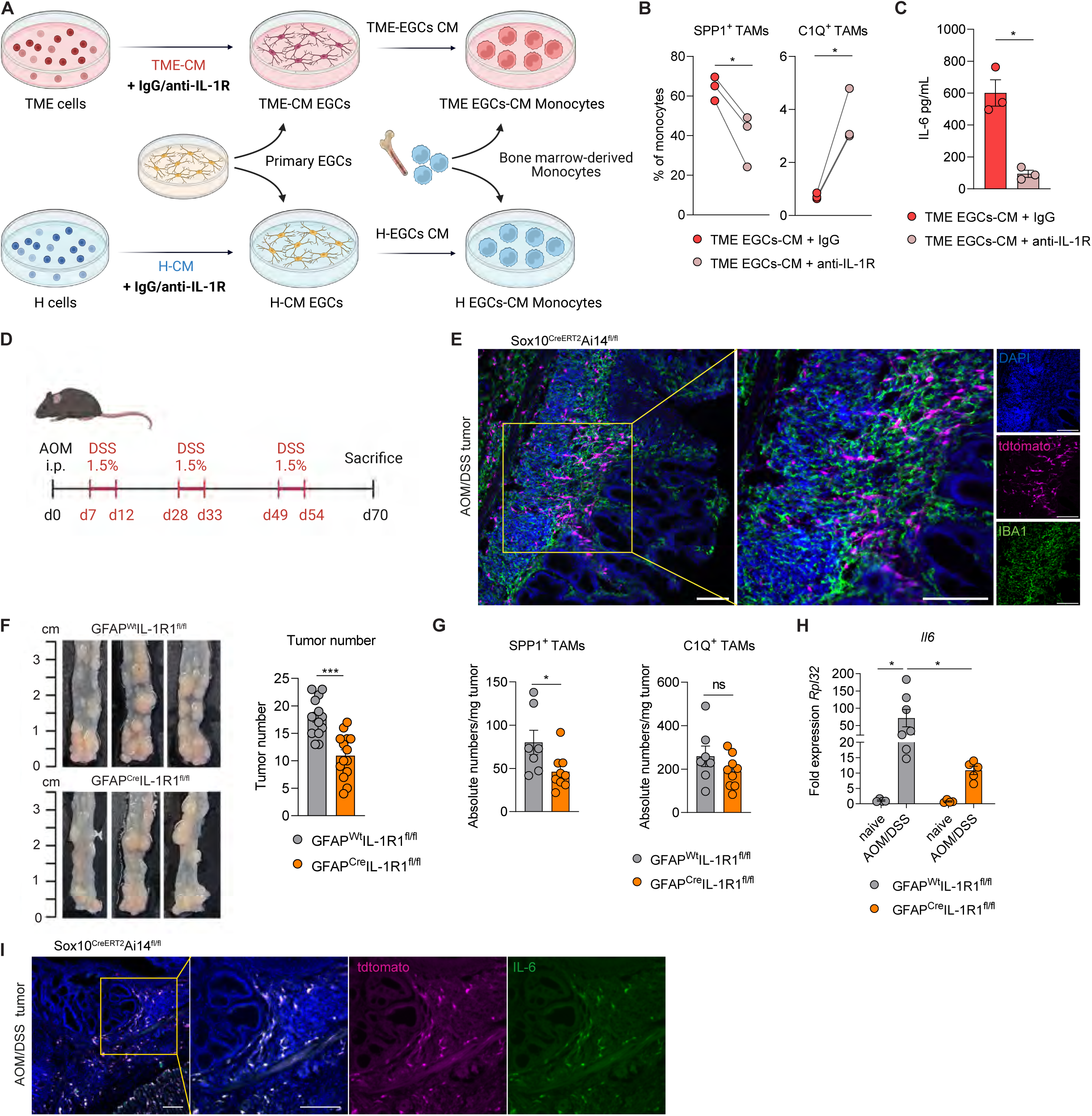
IL-1R signaling in EGCs promotes SPP1^+^ TAM differentiation and tumor progression. (A-C) Murine bone marrow-derived monocytes were cultured for 48h with supernatant of primary embryonic neurosphere-derived EGCs, which were pre-incubated for 24h with tumor microenvironment conditioned medium (TME-CM) together with isotype IgG or anti-IL-1R (5 µg/mL each) *(n = 3).* Experimental design (A). FACS quantification of SPP1^+^ TAMs and C1Q^+^ TAMs gated on monocytes after stimuli (B). IL-6 concentration in the conditioned medium of TME EGCs (C). (D) Schematic representation of experimental set-up of the murine AOM/DSS CRC model. Mice were subjected to an intraperitoneal (i.p.) injection with azoxymethane (AOM, 10 mg/kg body weight) at day(d)0. Starting from d7, mice underwent 3 repetitive cycles of 1.5% dextran sodium sulfate (DSS) (or 2% for Sox10^CreERT2^Ai14^fl/fl^ mice) in drinking water as indicated. (E) Sox10^CreERT2^Ai14^fl/fl^ mice were i.p. injected with Tamoxifen (1 mg in 100 µL sterile corn oil) on d-7, – 6, and –5. Subsequently, mice were subjected to the AOM/DSS model as described in Figure 6D. Representative image showing tdtomato (magenta), IBA1 (green), and DAPI (blue) in a tumor section at d70 (scale bar 100 µm). (F-H) GFAP^Wt^IL-1R1^fl/fl^ and GFAP^Cre^IL-1R1^fl/fl^ mice were subjected to the AOM/DSS model as described in Figure 6D. Tumor number, tumor tissue, and TAMs infiltration were collected or assessed at d70. Tumor numbers of GFAP^Wt^IL-1R1^fl/fl^ and GFAP^Cre^IL-1R1^fl/fl^ littermates with representative images (left) and quantitative comparison of the tumor numbers (right) *(n = 14)* (F). Corresponding absolute numbers of SPP1^+^ and C1Q^+^ TAMs per mg tumor tissue (*n = 7* GFAP^Wt^IL-1R1^fl/fl^, *n = 9* GFAP^Cre^IL-1R1^fl/fl^) (G). Relative mRNA levels for *Il6*, normalized to the housekeeping gene *Rpl32* in naive (*n = 4)* and AOM/DSS treated mice (*n = 7* GFAP^Wt^IL-1R1^fl/fl^, *n = 5* GFAP^Cre^IL-1R1^fl/fl^) (H). (I) Sox10^CreERT2^Ai14^fl/fl^ mice were subjected to the AOM/DSS model as described in Figure 6D. Representative image of tdtomato (magenta), IL-6 (green), and DAPI (blue) in tumor section at d70 (scale bar 100 µm). Data are represented as mean ± SEM (C, F-H). Statistical analysis: paired t-test for (B-C), unpaired t-test (F), unpaired Mann-Whitney test (G), and two-way ANOVA test with correction for multiple comparisons (H). *p < 0.05, *** p <0.0005, ns not significant. See also Figure S4 and Table S3.

To further analyze the role of IL-1R/IL-6 axis in this novel enteric glia and TAMs neuroimmune interaction *in vivo*, we utilized an inflammation-triggered CRC model induced by AOM/DSS (Figure 6D). Firstly, using the glia reporter mouse line Sox10^CreERT2^Ai14^fl/fl^, we confirmed spatial proximity of EGCs (tdtomato^+^) and TAMs (IBA1^+^) in the tumor regions as we previously observed in the orthotopic model (Figure 6E and Figure S4B). Transcriptomic analysis comparing AOM/DSS-induced tumors to naive tissue in wild-type mice additionally supports the involvement of the IL-1R/IL-6 pathway in EGC-TAM crosstalk within this model. In particular, transcriptomic differences highlighted an increase expression of CRC EGC markers, as well as TAM signature genes together with increase IL-1 signaling (Figure S4C and S4D).

To conclusively establish the role of IL-1-activated EGCs in *in vivo* CRC development, we subjected mice with a glial-specific knockout of IL-1R1 (GFAP^Cre^IL-1R1^fl/fl^) and their wild-type littermates (GFAP^Wt^IL-1R1^fl/fl^) to AOM/DSS (Figure S4E). Consistent with the role of enteric glia in the orthotopic CRC model, the number of colonic tumors was diminished in AOM/DSS-treated GFAP^Cre^IL-1R1^fl/fl^ compared to littermate GFAP^Wt^IL-1R1^fl/fl^ mice (Figure 6F), while no difference in weight loss were detectable between the two genotypes (Figure S4F). Interestingly, glial-specific IL-1R deficiency correlated with a decline in SPP1^+^ TAMs, but C1Q^+^ TAM levels remained unchanged (Figure 6G). Importantly, glial-specific IL-1R deficiency was further associated with a reduce *Il6* gene expression in tumor tissues of GFAP^Cre^IL-1R1^fl/fl^ compared to control GFAP^Wt^IL-1R1^fl/fl^ tumor lesions (Figure 6H). Consistently, IL-6 staining in tumor sections from the EGC reporter mouse line, Sox10^CreERT2^Ai14^fl/fl^, revealed that IL-6 protein co-localized with tdtomato^+^ glial cells in AOM/DSS tumors, confirming enteric glia as important source of IL-6 in the colonic TME (Figure 6I and S4G).

In conclusion, these results provide additional *in vivo* evidence highlighting the IL-1-dependent interplay between EGCs and SPP1^+^ TAMs in CRC.

### IL-1R induced-CRC EGC phenotype in patients affected by CRC

After identifying enteric glia-immune interactions in pre-clinical models of CRC, we investigated whether a similar process might influence disease progression in CRC patients. Initially, spatial tissue co-localization of EGCs and TAMs was confirmed in patient-derived CRC samples (Figure 7A). Next, the possible contribution of EGCs in disease outcome was defined using the colon and rectal cancer datasets from The Cancer Genome Atlas (TCGA-COAD and READ). Here, we observed that CRC patients with a higher enteric glia transcriptomic signature (Supplementary Methods Table 3), consisting of genes highly expressed by EGCs as in previously published CRC scRNA-seq datasets (Lee et al., 2020; Drokhlyansky et al., 2020; Kinchen et al., 2018), presented with a decreased overall survival probability compared to patient with low EGC signature (Figures 7B, S5A and S5B;). In-depth characterization of the patients with high EGC gene signature revealed 79% of this group belonging to the mesenchymal consensus molecular subtype 4 (CMS4) (Guinney et al., 2015), defined by the stromal invasion phenotype (Figure S5C). In comparison, only minor differences were identified when divided based on stage, microsatellite stability, or intrinsic CMS (iCMS) (Figure S5C). Interestingly, CRC patients with pronounced EGC involvement also exhibit higher expression of SPP1^+^ TAM signature genes (Figure 7C; Supplementary Methods Table 3).

**Figure 7.**
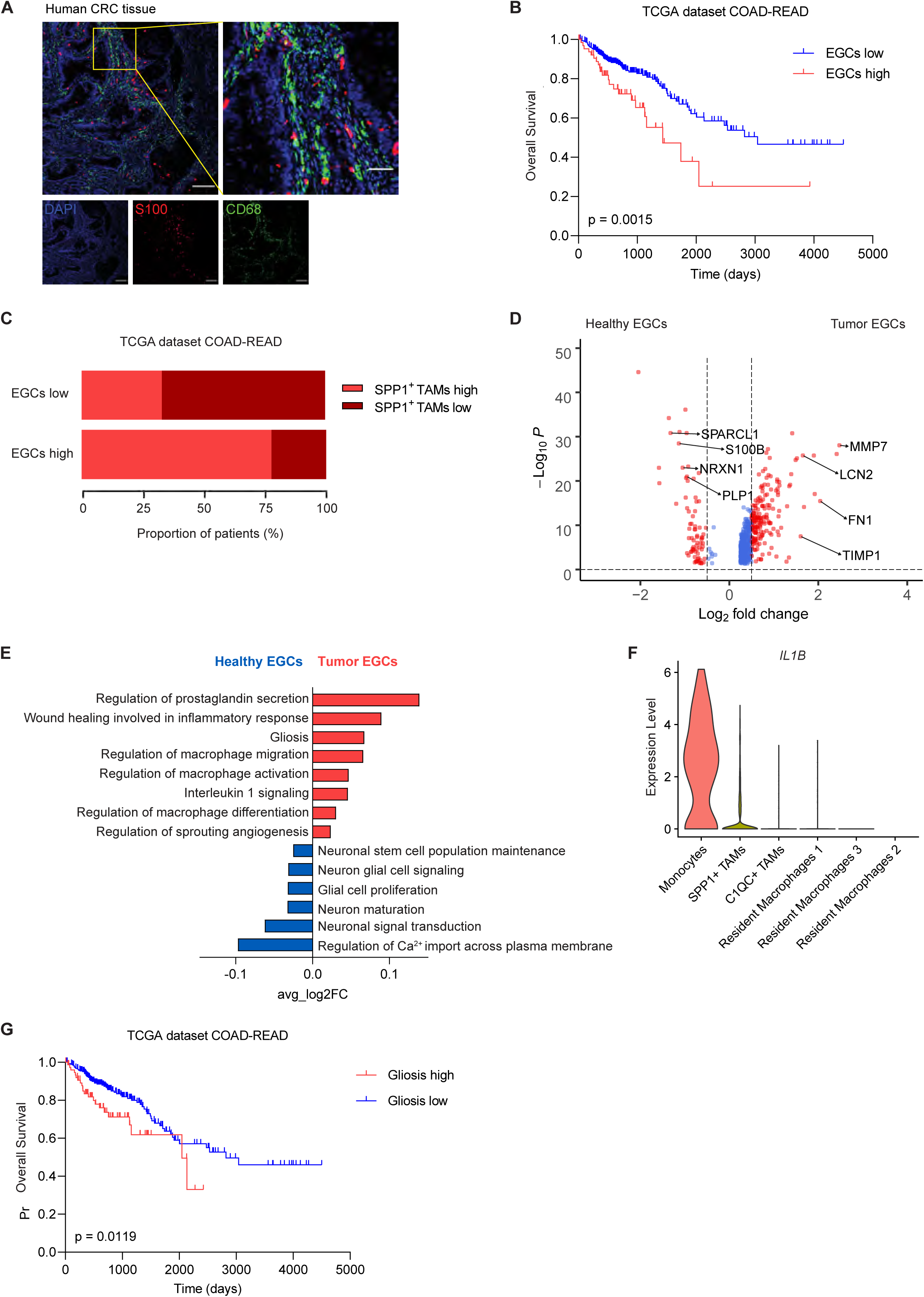
IL-1R induced-CRC EGC phenotype is conserved in patients affected by CRC. (A) Representative image showing S100 (red), CD68 (green) and DAPI (blue) in a human CRC tissue section (scale bar 200 µm and 50 µm). (B-C) TCGA COAD and READ patients stratified based on their expression of the EGCs signature genes. Heatmap of patients clustering (*n = 309* EGCs low, *n = 67* EGCs high). Kaplan-Meier overall survival curve for EGCs high and low patients (B). The proportion of EGCs high and low patients classified based on high or low SPP1^+^ TAMs gene signature expression (C). (D-E) Transcriptome analysis of tumor EGCs in CRC patients (KUL3 Dataset, Lee H. O. et al. 2020, *n = 5*). Volcano plot of differentially expressed genes between healthy and tumor EGCs (D), highlighting genes defining the tumor EGCs signature. Gene Set Enrichment Analysis presenting GO terms of interest (E). Violin plot showing expression of *IL1B* in the tumor-infiltrating myeloid cell clusters of human CRC in the KUL3 Dataset, Lee H. O. et al. 2020 (*n = 5*). Kaplan-Meier overall survival curve for TCGA dataset COAD-READ patients stratified based on their expression of the gliosis signature genes. (*n = 299* gliosis low, *n = 77* gliosis high). Statistical analysis: Mantel cox test (B, G). See also Figure S5 and Table S2.

Next, to assess if the murine CRC EGC transcriptional signature was conserved also in human CRC EGCs, we analyzed EGCs identified in a scRNA-seq dataset containing both CRC lesions and unaffected colonic tissues (Lee et al., 2020) (Figure S5D and S5E). By comparing gene expression profiles of tumor and healthy EGCs, we identified 589 genes specifically expressed in human CRC EGCs (Figures 7D; Table S2). Strikingly, among the top differentially expressed CRC EGC genes, we identified the two key murine CRC EGC marker genes *Lcn2* and *Timp1*. Using gene set enrichment analysis, we found that human EGC populations differentiate along the same homeostatic and tumor pathogenesis pathway transcriptomic signatures (Figure 7E) as seen in our murine EGCs (Figure 2E). Importantly, the significance of IL-1 signaling in the differentiation of patient CRC EGC was further confirmed by the increase transcriptomic signatures for “*Interleukin-1 signaling*” and “*Gliosis*” as well as “*Regulation of macrophage differentiation, activation and migration*” compared to healthy colonic EGCs (Figure 7E). In alignment with our pre-clinical findings, cell-population profiling of human CRC samples identified tumor-infiltrating monocytes as the primary source to IL-1β production (Figures 7F and S5F). Lastly, the activated tumor EGC state was also indicative of disease outcome, as patients with a more pronounced gliosis gene signature (Supplementary Methods Table 3) (Schneider et al., 2022), have a less favorable prognosis with decreased overall survival (Figure 7G). This indicates that both the presence of EGCs (Figure 7B) and their distinct IL-1 induced CRC EGC phenotype play roles in determining CRC prognosis.

## DISCUSSION

Using a variety of *in vitro* and *in vivo* models we uncover a previously unknown positive feedback loop between EGCs and TAMs in CRC. More specifically, we found that monocytes and monocyte-derived macrophages within the TME are the main producers of IL-1, inducing a pro-tumorigenic reactive phenotype in EGCs. IL-1 activated CRC EGCs via IL-6, in turn, directly promote the differentiation of tumor-infiltrating monocytes towards SPP1^+^ TAMs. Importantly, the unique reactive phenotype of tumor EGCs correlates with worse disease outcomes, as observed in both pre-clinical CRC mouse models and in patients with CRC. Here we provide new insights into the CRC pathogenesis, uncovering a previously unidentified neuroimmune interaction between EGCs and TAMs.

EGCs are a highly plastic cells that can rapidly adapt their functions under the influence of microenvironmental cues (Seguella and Gulbransen, 2021). Recent studies have identified specific immunomodulatory factors, including IFN-γ, IL-1, and ATP, as triggers of enteric glia phenotypic and functional reprogramming in both homeostasis and diseased conditions (Progatzky et al., 2021; Schneider et al., 2022, 2021). In particular, IL-1-mediated EGC reactivity and its effects on immune cell modulation have been extensively studied in the context of intestinal inflammation (Schneider et al., 2022; Stoffels et al., 2014). However, the mechanisms underlying these processes in CRC were not yet understood. Using an unbiased approach combining murine bulk and human scRNA-seq, our investigation pinpointed TME-derived IL-1 as the principal initiator of EGC reactivity. Furthermore, we found that this IL-1-triggered EGC activation coincides with a profoundly immunomodulatory transcriptional signature in CRC EGCs. Of note, IL-1R signaling in EGCs may hold relevance for additional functions of CRC EGCs, as *in vitro* studies indicated the significance of IL-1 in EGC-cancer stem cell interactions (Valès et al., 2019). Interestingly, we identified tumor-infiltrating monocytes and macrophages as the main source of IL-1 within the tumor. Nevertheless, although our *in vivo* data and scRNAseq studies could not verify epithelial cells as a significant IL-1 source in CRC, low amounts of tumor epithelial IL-1 might also contribute to EGC activation during CRC (Valès et al., 2019).

Our study utilizes single-cell and bulk RNA-seq techniques to better understand and predict the interactions between EGCs and TAMs within the colonic TME. In our murine orthotopic CRC model, we identified two distinct subsets of TAMs with different ontogeny and properties. The C1Q^+^ TAMs, which preferentially express genes involved in phagocytosis and antigen presentation, coexist in the TME with SPP1^+^ TAMs that are enriched for factors regulating angiogenesis and extracellular matrix, suggesting their key role in colon tumorigenesis. This dichotomy recently also identified in patients with colorectal cancer (Zhang et al., 2020), supports the relevance of our findings to human disease. Importantly, our findings underscore a novel EGC-TAM interaction, showing that IL-1 activated EGCs drive monocyte differentiation into pro-tumorigenic SPP1^+^ macrophages through IL-6. Thereby, our study identified EGCs as an additional important regulator of SPP1^+^ TAM differentiation that, together with other cancer-associated stromal cells, may contribute to tumor progression (Qi et al., 2022; Luo et al., 2022). Consistent with this, pan-cancer transcriptome analysis has pinpointed SPP1^+^ TAMs as the most pro-tumorigenic macrophage subset across various cancers, including CRC (Cheng et al., 2021). Hence, inhibiting the differentiation of SPP1^+^ TAMs by blocking IL-1R signaling in EGCs may significantly impede tumor progression.

Furthermore, our data also point to a functional associated between CRC EGCs and monocyte migration, as reflected by increased *Ccl2* expression in CRC glia. Considering that the tumor monocyte population decreased upon EGC depletion, we speculate that CRC EGC-derived chemokines (i.e., CCL2 and CXCL5) could also promote the infiltration of monocytes in the colonic tumor site. This would be in line with our recent findings showing early expression of CCL2 by reactive EGCs in the context of intestinal inflammation (Stakenborg et al., 2022). However, further research will need to determine whether the pro-tumorigenic role of EGCs is exerted solely on the SPP1^+^ TAMs or whether EGCs also affect other immune or stromal cells via glial-derived factors. Recent work by Progatzky et al. provides supportive evidence for this, demonstrating that EGCs are involved in a protective immune and stromal response to control parasitic insult in the gut (Progatzky et al., 2021).

Consistent with the identification of a pro-tumorigenic EGC phenotype in mice, we found that also in patients the reactive EGC transcriptomic signature was associated with reduced overall survival. Therefore, we could speculate that our newly identified CRC EGC gene signature might be used as a potential biomarker to predict disease outcomes. In line with the stromal nature of EGCs, we demonstrated that the vast majority of patients with high EGC involvement belonged to the consensus molecular subtype 4 (CMS4), which is characterized by a mesenchymal-like phenotype, a strong stromal infiltration and the worst overall and relapse-free survival compared to the other CMS subtypes (Guinney et al., 2015). Gene ontology analysis revealed that also the human tumor glial cells were enriched for immunomodulatory transcriptional programs related to macrophage differentiation, leading to the assumption that CRC EGC-derived signals modulate TAMs also in patients. In line, human tumor EGCs also displayed enriched GO terms for “Interleukin-1 signalling” and “Gliosis”, hinting at a similar EGC activation in CRC as in our preclinical models. Moreover, various studies demonstrated increased IL-6 levels in both tumor tissues and serum samples of human CRC patients compared to healthy controls (Komoda et al., 1998; Galizia et al., 2002). Consistently, immune-related pathways, including IL-6 signalling, were enriched in CRC EGCs in a human single-cell data set published by Qi et al. (Qi et al., 2022).

Our research elucidates the role of the IL-1/IL-6 axis in glial-immune communication in CRC, which could potentially be of relevance to various other tumors exhibiting neuronal infiltration, a features often associated with less favorable disease outcomes (Pundavela et al., 2015; Renz et al., 2018; Hayakawa et al., 2017; Albergotti et al., 2017; Zhu et al., 2018; Murakami et al., 2013; Magnon et al., 2013). Apart from EGCs, peripheral glial cells, including Schwann cells, are known to play a crucial role in cancer pathophysiology, as demonstrated in pancreatic ductal adenocarcinoma, lung cancer, and melanoma (Deborde et al., 2022; Zhou et al., 2020; Shurin et al., 2019). Consistent with the pro-tumorigenic functions of EGCs in CRC, studies in melanoma models have shown that tumor Schwann cells favor the differentiation of pro-tumorigenic macrophages enhancing tumor growth (Shurin et al., 2019). Overall, glial-immune crosstalk might be an overlooked critical component of tumor pathophysiology in many cancer types beyond CRC.

In conclusion, our study reveals a critical role for IL-1R signaling in driving enteric glia-macrophage interactions in CRC pathogenesis. Our research provides essential insight into the complex neuroimmune mechanisms underlying the development of this disease, shedding light on potential novel biomarkers and specific therapeutic targets that hold the promise of transforming the management of this devastating disease.

## ACKNOWLEDGMENTS

We acknowledge all members of Prof. Matteoli’s laboratory and Prof. Wehneŕs laboratory for the support and scientific discussions. We would like to thank Tine Gomers, Karlien Vranken and Renata Siqueira de Mello (TARGID, KU Leuven) and Patrik Efferz and Bianca Schneiker (Department of Surgery, University Hospital Bonn) for technical assistance during experiments. Furthermore, we would like to thank Ally Peddle and Yourae Hong (Molecular Digestive Oncology, Department, KU Leuven), Lukas Ferreira Maciel (Laboratory for Molecular Cancer Biology, VIB-KU Leuven) and Florent Petitprez (MRC Centre for Reproductive Health, University of Edinburgh) for their scientific support on bio-informatic analysis. Within KU Leuven we would like to acknowledge the following core facilities: FACS Core, Genomics Core (UZ Leuven), LiMoNe VIB Bioimaging Core and VIB Center for Brain & Disease Research. We would like to thank the Cell and Tissue Imaging Cluster (KU Leuven) for the usage of the Zeiss LSM 880 – Airyscan (supported by Hercules AKUL/15/ 37_GOH1816N and FWO G.0929.15 to Prof. Pieter Vanden Berghe). We would like to thank the support from the Core Facilities of the Medical Faculty, University of Bonn, specifically, the Analytical Proteomics Core, funded by the Deutsche Forschungsgemeinschaft (DFG) – project 386936527, the Bioinformatics Data Analysis Core, the Microscopy Core Facility funded by the DFG – project 266686698, and the Next Generation Sequencing Core. We thank Dr. Vassilis Pachnis for sharing the Sox10^CreERT2^ mice with us. BioRender was used for creating graphical images.

V.D.S. was supported by a Stichting tegen Kanker postdoctoral fellowship. S.S. was supported by KU Leuven-University of Melbourne Global PhD (GPUM/22/020). F.B. was supported by the KU Leuven-University of Edinburgh Global PhD (GPUE/20/003). B.K. was supported by the Taiwan – KU Leuven PhD Scholarship. M.V. was supported by a Fonds voor Wetenschappelijk Onderzoek Vlaanderen (FWO, 11L0822N) PhD fellowship. S.I. was supported by a MSCA-IF (79756–GLIAMAC) and a fellowship from the European Crohn’s and Colitis Organization (ECCO). S.V. and S.T. were supported by the FWO grant G067821N and Stichting tegen Kanker grant F/2020/1512. G.M.’s lab was supported by FWO grants G0D8317N, G0A7919N, G086721N, G088816N and S008419N, KU Leuven Internal Funds (C12/15/016 and C14/17/097). S.W. and L.S. were supported by the DFG-funded Immunosensation^2^ cluster of excellence EXC2151-190873048. S.W. and R.S. received funding from BONFOR. B.G.R. received funding from the Deutsche Krebshilfe through a Mildred Scheel Nachwuchszentrum Grant (70113307).

## AUTHOR CONTRIBUTIONS

Conceptualization, L.V.B., V.D.S., L.S., R.S., S.W. and G.M.; Methodology, L.V.B., V.D.S., L.S., S.W. and G.M.; Software, S.S., S.A., Z.H., J.H., L.V.B. and L.S.; Validation, L.V.B., V.D.S. and L.S.; Formal Analysis, L.V.B., V.D.S., L.S., S.S. and S.A.; Investigation, L.V.B., V.D.S., L.S., S.S., S.A., F.B., L.Z., S.V., B.K., B.G.R., M.T., M.V. and S.I.; Resources, G.M., S.W., S.T. and F.D.S.; Data Curation, L.V.B., V.D.S., L.S., S.S., S.A., S.V., Z.H. and J.H.; Writing – original draft, L.V.B., V.D.S. and L.S.; Writing – review & editing, all; Visualization, L.V.B., V.D.S., L.S., S.S., S.A., and Z.H.; Supervision, G.M., S.W., S.T., F.D.S., R.S., M.S. and V.D.S.; Project administration, L.V.B., V.D.S., L.S., R.S., S.W. and G.M.; Funding acquisition, V.D.S., S.I., G.M. and S.W..

## DECLARATION OF INTERESTS

The authors declare no competing interests.

## INCLUSION AND DIVERSITY

We support inclusive, diverse, and equitable conduct of research.

## Notes

### Competing Interest Statement

The authors have declared no competing interest.

### Summary of Updates

Updated Manuscript

