## Supplemental Methods and Figures for "IL-1R signaling drives enteric glia-macrophage interactions in colorectal cancer"

**SUPPLEMENTARY MATERIALS & METHODS**

**Materials availability.** This study did not generate new unique reagents.

**MC38 cell line**. Murine colon adenocarcinoma cell line MC38 (NCI, ENH204-FP) was kindly provided by Prof. Max Mazzone (VIB - KU Leuven). The MC38 cell line was maintained in 5% $CO_2$ at 37°C in high glucose Dulbecco's Modified Eagle Medium (DMEM) (Gibco, ThermoFisher Scientific) supplemented with 10% Fetal Bovine Serum (FBS) (Biowest), 100 µg/mL Penicillin and Streptomycin, 2 mM L-glutamine, 10 mM N-2-hydroxyethylpiperazine-N-2-ethane sulfonic acid (HEPES), 1 mM sodium pyruvate, 50 µM 2-Mercaptoethanol and 1X Non-Essential Amino Acids (all from Gibco, ThermoFisher Scientific).

**Embryonic neurosphere-derived enteric glial cells (EGC) culture.** Embryonic neurosphere-derived EGCs were obtained as previously described (Stakenborg et al., 2022). Briefly, total intestines from E13.5 C57BL/6J mice were digested with collagenase D (0.5 mg/mL; Roche) and DNase I (0.1 mg/mL; Roche) in DMEM/F-12 (Gibco, ThermoFisher Scientific) for 1 hour at 37°C under gentle agitation. After digestion, tissue was filtered through a 70 µm cell strainer and cells were cultured in a $CO_2$ incubator at 37°C in DMEM/F-12, 100 µg/mL Penicillin and Streptomycin, 2 mM L-glutamine, 10 mM HEPES, 1 mM sodium pyruvate, 50 µM 2-Mercaptoethanol supplemented with 1x B27 (Gibco, ThermoFisher Scientific), 40 ng/mL Epidermal Growth Factor (EGF) (Stemcell Technologies) and 20 ng/mL Fibroblast Growth Factors (FGF) (Invitrogen, ThermoFisher Scientific). After a minimum of 1 week of culture, neurospheres were treated with NeuroCult™ Chemical Dissociation Kit (Stemcell Technologies) according to the manufacturer's protocol and filtered through a 70-µm cell strainer. Cells were seeded on Poly-D-Lysine (PDL solution, 1.0 mg/mL, Sigma Aldrich) coated plates and differentiated in DMEM medium supplemented with 10% FBS, 100 µg/mL Penicillin and Streptomycin, 2 mM L- glutamine, 10 mM HEPES, 1 mM sodium pyruvate, 50 µM 2-Mercaptoethanol (DMEM complete medium) until confluence for 5 days to obtain primary EGCs. For IL-1 stimulation experiments, $5 \times 10^4$ neurosphere-derived EGCs/mL were stimulated with or without recombinant murine 10 ng/mL IL-1α (Peptrotech) and/or 10 ng/mL IL-1β (Sigma Aldrich) for 24 hours.

**Neurosphere-derived adult enteric glial cells (EGC) culture.** Neurosphere-derived adult EGC cultures were obtained as described previously (Schneider et al., 2022). Briefly, small intestines of 8-16 weeks old C57BL/6 mice were harvested, cleansed, cut in 3-5 cm long segments and kept in oxygenated Krebs-Henseleit buffer (126 mM NaCl; 2.5 mM KCl: 25 mM $NaHCO_3$; 1.2 mM $NaH_2PO_4$; 1.2 mM $MgCl_2$; 2.5 mM $CaCl_2$, 100 IU/mL Penicillin, 100 IU/mL Streptomycin and 2.5 µg/mL Amphotericin). For each segment, the muscularis layer was peeled and collected for digestion. Muscularis tissues were incubated for 15 min in DMEM containing Protease Type 1 (0.25 mg/mL, Sigma-Aldrich) and Collagenase A (1 mg/mL, Sigma-Aldrich) at 37 °C and 150 rpm. Digestion was stopped with DMEM containing 10% FBS (Sigma-Aldrich) and cells were cultured in proliferation medium (neurobasal medium with 100 IU/ Penicillin, 100 µg/mL Streptomycin, 2.5 µg/mL Amphotericin (all ThermoFisher Scientific), FGF and EGF (both 20 ng/mL, Immunotools) at 37°C, 5% $CO_2$ to promote neurosphere formation. After 1 week in culture, enteric neurospheres were dissociated with trypsin (0.25%, ThermoFisher Scientific) for 5 min at 37 °C and differentiated at 50% confluency on Matrigel (100 µg/mL, Corning) coated 6 well plates for 1 week in differentiation medium (neurobasal medium with 100 IU/ Penicillin, 100 µg/mL Streptomycin, 2.5 µg/mL Amphotericin, B27, N2 (all Thermo Scientific) and EGF (2 ng/mL, Immunotools). For liquid chromatography/mass spectrometry (LC/MS) experiments, mature EGCs were treated with or without IL-1β (10 ng/mL, Immunotools) for 24 hours. Conditioned media were collected and concentrated using Pierce™ Protein Concentrators, 3K MWCO (ThermoFisher Scientific) according to the manufacturer's instructions. After denaturation at 95°C for 5 min, samples were snap-frozen and kept at -80°C until further processing.

**Bone marrow-derived monocyte isolation and stimulation**. Murine bone marrow (BM)-derived monocytes were isolated from C57BL/6 mice. Briefly, the tibia and femur were dissected, and BM cells were flushed with DMEM high glucose supplemented with 10% FBS. After cells were collected and counted, monocytes were isolated with the EasySepTM Mouse monocyte isolation kit (Stemcell Technologies) according to the manufacturer's instructions. Next, 5 x $10^5$ monocytes were stimulated for 48 hours with 1 mL of H-CM, TME-CM, or with the supernatant of *in vitro* embryonic neurosphere-derived EGCs pre-incubated for 24h with H-CM or TME-CM. For IL-1R blocking experiments, 5 µg/mL isotype IgG (BioXCell) or 5 µg/mL anti-IL-1R (BioXCell) was removed from the EGCs supernatant by using 20 µL/mL Dynabeads™ Protein G (ThermoFisher Scientific) according to the manufacturer's instructions. For IL-6 neutralization experiments, 20 µL/mL Dynabeads™ Protein G (ThermoFisher Scientific) together with 5 µg/mL anti-IL-6 (R&D Systems) or 5 µg/mL isotype IgG (R&D Systems) was added to the supernatant of H-CM EGCs and TME-CM EGCs and incubated for 2 hours before removal by taking advantage of magnetisation with the DynaMag™-2 Magnet (ThemoFisher Scientific).

**AOM-DSS model.** For Sox10$^{CreERT2}$Ai14$^{fl/fl}$ mice subjected to the AOM/DSS model, slight adjustments were made to the protocol. Female Sox10$^{CreERT2}$Ai14$^{fl/fl}$ mice were i.p. injected with Tamoxifen (MP biomedicals, 1 mg in 100 µL sterile corn oil) on days -7, -6, and -5. On day 0, the AOM/DSS model was started as described above, but here mice were subjected to a DSS concentration of 2% in drinking water since this strain was less susceptible to DSS. On day 70, colons were harvested and cryo-embedded as swiss rolls for immunohistochemistry.

**MILAN multiplex immunohistochemistry of tumor tissue sections of patients with CRC.** Multiplex immunohistochemistry and analysis were performed according to a previously published method

((Pombo Antunes et al., 2021; Maria Bosisio et al., 2020; Naulaerts et al., 2023) and https://doi.org/10.21203/rs.2.1646/v5). Briefly, tissue sections (3 µm thickness) were prepared from formalin-fixed paraffin-embedded human CRC samples (collected at the UZ/KU Leuven biobank according to protocol S66460). First, FFPE-tissue slides were deparaffinized by sequentially placing them in xylene, 100% ethanol and 70% ethanol. Following dewaxing, antigen retrieval was performed using PT link (Agilent) using 10 mM Ethylenediaminetetraacetic acid (EDTA) in Tris-buffer pH 8. Immunofluorescence staining was performed using Bond RX Fully Automated Research Stainer (Leica Biosystems) with the primary antibodies rabbit anti-S100B (Dako) and mouse anti-CD68 (Invitrogen, ThermoFisher Scientific). The sections were incubated for 4h with the primary antibodies, washed several times and afterwards stained for 30 min with fluorescently labelled secondary antibodies (Alexa fluor 647 donkey anti-rabbit and Alexa Fluor 488 goat anti-mouse respectively). Slides were then incubated for 10 min with a buffer containing 4,6-diamidino-2-phenylindole (DAPI), after which mounting medium (50% glycerol; 584mM C12H22O11; 10mM Phosphate, 154mM NaCl; pH 7,5) and a coverslip (Agilent, ref. CR12230-7) were manually applied to the slides. Then the slides were scanned using a Zeiss Axio Scan Z.1 (Zeiss) at 10x magnification. Utilize ImageJ (1.53T) and Qu path (0.3.2) were used for the region selection and to subtract background and tissue autofluorescence. Further analysis was performed by using ImageJ.

**Immunohistochemistry and Immunofluorescence.** Orthotopic tumors were fixed overnight (ON) at 4°C in Periodate-Lysine-Paraformaldehyde (PLP) buffer consisting of Milli-Q Water supplemented with 1% paraformaldehyde, 0.075 M lysine (pH 7.4), 0.037 M sodium phosphate (pH 7.4) and 0.01 M $NaIO_4$ (all from Sigma-Aldrich). Samples were washed three times with Milli-Q Water supplemented with 0.037 M sodium phosphate (pH 7.4), followed by a minimum of 4h incubation in 30% sucrose (VWR chemicals) in PBS. Then, samples were embedded in OCT (Scigen) and stored at -80°C until usage.

Preceding immunohistochemical staining, 7-µm orthotopic tumor tissue sections on SuperFrost Plus™ Adhesion slides (Epredia) were exposed to two washes with HistoChoice Cleaning Agent for 2 min each (Sigma-Aldrich) and subsequent hydration with Ethanol 100% for 2 min each (Merck) followed by deionized water. Then haematoxylin and eosin (both from Leica) staining was performed using standard procedures. Imaging was performed with Nikon Marzhauser Slide Express2, processed and analysed using ImageJ.

Preceding immunofluorescent staining, tissues were sectioned to 7-µm thickness on SuperFrost Plus™ Adhesion slides (Epredia) and blocked with blocking buffer (PBS containing 0.02% Sodium azide (Sigma-Aldrich), 0.3% donkey serum (Jackson), and 3% Bovine Serum Albumin (BSA, Serva) for 2h at room temperature (RT). Subsequently, samples were incubated ON at 4°C with the following primary antibodies: 1:500 rat anti-F4/80, BioRad and/or 1:300 rabbit anti-GFAP Dako in staining buffer (blocking buffer supplemented with 0.3% Triton X-100 (ThermoFisher, Scientific)). Then, samples were washed in PBS and incubated with DAPI (4′,6-Diamidine-2′-phenylindole dihydrochloride; Sigma-Aldrich) combined with the secondary antibodies: 1:1000 donkey anti-rat AF488 (Invitrogen, ThermoFisher Scientific), and/or 1:400 donkey anti-rabbit Cy5 (Jackson) in staining buffer for 2h at RT. Finally, samples were rinsed three times in PBS and mounted with SlowFade Diamond Antifade mounting (Invitrogen, ThermoFisher Scientific). Imaging was performed on the ZEISS LSM 880 confocal microscope and the pictures were analysed using ImageJ.

For immunofluorescent staining of AOM/DSS or naive swiss rolls of Sox10$^{CreERT2}$Ai14$^{fl/fl}$ mice were fixed ON at 4°C in 4% PFA. Samples were washed once with PBS followed by ON incubation at 4°C in 30% sucrose (Sigma) in PBS. Subsequently, swiss rolls were embedded in Tissue-Tek® O.C.T.™ Compound (Sakura) and stored at -80°C until usage.

Prior to immunofluorescent staining, AOM/DSS or naive Sox10$^{CreERT2}$Ai14$^{fl/fl}$ samples were sectioned to 14 µm thickness on SuperFrost Plus™ Adhesion slides (Epredia). Slides were washed three times in PBS and blocked with blocking buffer (PBS containing 3% donkey serum and 0.1% Triton X-100) for 1h at RT. Subsequently, primary antibody staining was performed ON at 4°C in staining buffer (blocking buffer diluted with PBS in a 1:1 ratio) using the following antibodies: rabbit anti-IBA1 (1:400, Abcam), rabbit anti-IL-6 (1:100, Abcam), or chicken anti-GFAP (1:1000, Biolegend). For IgG control staining, rabbit IgG (Dianova) was used in the same antibody concentration as rabbit anti-IL6. Slides were washed three times in PBS and secondary antibody staining was performed for 2h at RT with donkey anti-rabbit FITC (1:800, Dianova) or donkey anti-chicken FITC (1:800, Jackson) in staining buffer. After three more washes with PBS, slides were incubated with DAPI (Sigma-Aldrich) for 5 min at RT. After a final wash with PBS, slides were mounted with Shandon™ Immu-Mount™ (Epredia). Imaging was performed on the Nikon ECLIPSE Ti2 microscope using NIS-Elements AR software (version 5.41.01) or a Leica SP8 with LAS AF v3.x software for confocal images. Pictures were analysed using ImageJ.

For whole mount samples of GFAP$^{Cre}$Ai14$^{fl/fl}$ mice, the terminal colon was opened longitudinally, fixed with 4% PFA for 20 min, and washed with Krebs-henseleit buffer. *Muscularis externa* was peeled off the colonic tissue and permeabilized (1% Triton X-100 in PBS) for 20 min followed by blocking with blocking buffer (PBS containing 3% donkey serum and 0.1% Triton X-100) for 1h at RT. Antibody staining was performed as described above (for Sox10$^{CreERT2}$Ai14$^{fl/fl}$ samples) using chicken anti-GFAP (1:1000, Biolegend) and goat anti-SOX10 (1:1000, self-made) primary antibodies and donkey anti-chicken FITC (1:800, Jackson) and donkey anti-goat Cy5 (1:800, Sigma). Imaging was performed on a Leica SP8 with LAS AF v3.x software for confocal images. Pictures were analysed using ImageJ.

**Enzyme-linked Immunosorbent Assay (ELISA).** Healthy and tumor conditioned medium EGCs supernatants were collected and analysed for IL-6 and IL-1β content using sensitive commercial ELISA kits (R&D Systems, Minneapolis, MN and V-Plex Pro-inflammatory panel Meso Scale Discovery; MSD respectively) according to the manufacturer's instructions. The data were analysed with the Discovery Workbench 4.0 software (MSD).

**Western Blot.** Total proteins were extracted from mouse colonic tissues in T-PER buffer (ThermoFisher, Scientific) supplemented with 1 mM dithiothreitol, 10 mg/mL aprotinin, 10 mg/mL leupeptin, 1 mM phenylmethylsulfonyl fluoride, 1 mM Na3VO4 and 1 mM NaF (all from ThermoFisher, Scientific), by homogenization for 1 minute at 30 Hz (TissueLyser II, Qiagen). Lysates were clarified by centrifugation at 4 °C, 12000 g for 30 min and separated on sodium dodecyl sulphate (SDS)-polyacrylamide gel electrophoresis. Blot was incubated with the GFAP antibody (1:500 final concentration, Cell Signaling) followed by a secondary antibody conjugated to horseradish peroxidase (1:5000 final dilution; both from Dako Agilent Technologies). To ascertain equivalent loading of the lanes, the blot was stripped and incubated with an anti-vinculin antibody (1:5000 final dilution, Sigma-Aldrich). Computer-assisted scanning densitometry (GE Healthcare ImageQuant LAS 4000 Luminscent Image Analyzer) was used to analyze the intensity of the immunoreactive bands.

**Liquid Chromatography/Mass Spectrometry (LC/MS).** LC/MS analysis of adult neurosphere-derived EGC supernatants treated with or without IL-1β was performed by the Core Facility Analytical Proteomics of the University of Bonn as described in the following. All chemicals from Sigma unless otherwise noted. For LC/MS sample preparation, 70 µg of protein per sample was subjected to in-solution preparation of peptides with the iST-NHS 96x sample preparation kit (Preomics GmbH, Martinsried, Germany) according to the manufacturer's recommendations. 0.4 mg TMT10plex isobaric Mass Tag Labeling reagent (Thermo Scientific) was added to each sample and incubated at room temperature for 1 hour. 10 µL 5% hydroxylamine was used to quench the reaction. The preparation procedure was continued according to the iST-NHS kit instructions. Peptide concentration was determined with a colorimetric peptide assay (Thermo Scientific). Equal amounts of peptides were pooled and dried in a vacuum concentrator, dissolved in 20 mM ammonium formate (pH 10) and fractionated by reversed phase chromatography at elevated pH with a Reprosil 100 C18 column (3 µm 125 x 4 mm, Dr. Maisch GmbH, Ammerbuch-Entringen, Germany). 60 fractions were combined into 6 pools and dried in a vacuum concentrator.

Before measurement, peptides were re-dissolved in 0.1% formic acid (FA) to yield a 1 g/L solution and separated on a Dionex Ultimate 3000 RSLC nano HPLC system (Dionex GmbH, Idstein, Germany). The autosampler was operated in µL-pickup mode. 1 µL was injected onto a C18 analytical column (self-packed 400 mm length, 75 µm inner diameter, ReproSil-Pur 120 C18-AQ, 1.9 µm, Dr. Maisch). Peptides were separated during a linear gradient from 5% to 35% solvent B (90% acetonitrile, 0.1% FA) at 300 nL/min during 150 min. The nano-HPLC was coupled online to an Orbitrap Fusion Lumos Mass Spectrometer (Thermo Fisher Scientific, Bremen, Germany). Peptide ions between 330 and 1600 m/z were scanned in the Orbitrap detector every three seconds with a resolution of 120,000 (maximum fill time 50 ms, AGC target 100%). From MS3-based quantification, peptides were subjected either to collision induced dissociation for identification (CID: 0.7 Da isolation, normalized energy 30%) and fragments analyzed in the linear ion trap with AGC target 50% and a maximum fill time 35 ms, rapid mode. Fragmented peptide ions were excluded from repeat analysis for 30 s. The top 10 fragment ions were chosen for synchronous precursor selection and fragmented with higher energy CID (HCD: 3 Da MS2 isolation, 65% collision energy) for detection of reporter ions in the Orbitrap analyzer (range 100-180 m/z, resolution 50,000, maximum fill time 86 ms, AGC target 200%). Alternatively, peptides were only fragmented by HCD and fragment ions and reporter ions analyzed in the same spectrum (Orbitrap resolution 50,000).

Raw data processing and database search were performed with Proteome Discoverer software 2.5.0.400 (Thermo Fisher Scientific). Peptide identification was done with an in-house Mascot server version 2.8.1 (Matrix Science Ltd, London, UK). LC/MS data were searched against the Uniprot reference proteome mouse database (2022/05, 63628 sequences) and contaminants database (cRAP1) (Mellacheruvu et al., 2013). Precursor Ion m/z tolerance was 10 ppm, fragment ion tolerance 0.5 Da (CID). Tryptic peptides with up to two missed cleavages were searched. C6H11NO-modification of cysteines (delta mass of 113.08406) and TMT10plex on N-termini and lysines were set as static modifications. Oxidation was allowed as dynamic modification of methionine. Mascot results were evaluated by the Percolator algorithm version 3.02.12 as implemented in Proteome Discoverer (The et al., 2016). Spectra with identifications above 1% q-value were sent to a second round of database search with semi tryptic enzyme specificity (one missed cleavage allowed). Protein N-terminal acetylation, methionine oxidation, TMT10plex, and cysteine alkylation were then set as dynamic modifications. Actual FDR values were 0.2% (peptide spectrum matches) and 0.9% (peptides). Reporter ion intensities (most confident centroid) were extracted from the MS3 level, with SPS mass match >65%.

The statistical analyses of the peptide-spectrum match (PSM) level data were done by the Core Unit for Bioinformatics Data Analysis of the University of Bonn. Analyses were carried out in R environment (R version 4.2) using an in-house developed workflow. Non-unique peptides and single-hit proteins (proteins identified/quantified by only one peptide) were filtered-out prior to the statistical analysis.

From all available fractions, only those with the least number of missing values per feature and maximum average intensity across all TMT labels were selected. The PSM-level data were then log-transformed and scaled such that all the samples have the same median values (median normalization method). Next, the normalized data was aggregated to protein-level by applying the Tukey's median polish method. The statistical analysis was performed using the R package limma (Ritchie et al., 2015). For each statistical contrast, the resulting P-values were adjusted for multiple testing. The false discovery rates (FDR) were calculated by the Benjamini-Hochberg method.

**Isolation of tumor-infiltrating leukocytes.** Tumor-bearing mice were sacrificed at the described time points. After peeling off the muscularis layer from the orthotopic tumors, tissues were first cut in 1 mm pieces, and then went under mechanical and enzymatic digestion for 30 min in DMEM with 2.5% FBS, 100 µg/mL Penicillin and Streptomycin, 200 U/mL collagenase IV (Gibco, ThermoFisher Scientific) and 125 µg/mL type II dispase (Gibco, ThermoFisher Scientific). AOM/DSS induced tumors and healthy colon samples were peeled off the muscularis layer and underwent epithelial removal by vigorous shaking in Hanks' balanced salt solution (HBSS) with phenol red (Gibco, ThermoFisher Scientific) containing 1% FBS, 100 µg/mL Penicillin and Streptomycin, 1 mM EDTA (Invitrogen, ThermoFisher Scientific) and 1 mM dithiothreitol (DTT) (Sigma- Aldrich) for 8 min at 37°C. A second incubation step was performed for 8 min at 37°C in the same medium without DTT. After washing in wash medium (DMEM with 2.5% FBS and 100 µg/mL Penicillin and Streptomycin), the remaining tissue was cut into small pieces and digested for 30 min at 37°C in pre-warmed alpha Minimum Essential Medium (MEM) (Lonza) containing 5% FBS, 100 µg/mL Penicillin and Streptomycin, 5 U/mL DNase (Roche), 1 mg/mL dispase (Gibco, ThermoFisher Scientific), 1.25 mg/mL Collagenase D (Roche) and 0.85 mg/mL Collagenase V (Sigma-Aldrich). Independent of tumor origin, cells were then filtered through a 70-µm cell strainer (BD Falcon), washed with PBS, and stained with fluorophore-conjugated antibodies.

**FACS staining and analysis.** Single-cell suspensions (obtained as described above) were incubated for 15 min with mouse FcR Blocking Reagent (1:100 BD Pharmingen) at 4 °C. Next, cells were stained for surface markers (see Methods Table 1 for antibodies list) and incubated for 20 min incubation at 4 °C, then cells were washed with FACS buffer (0.5% FBS and 2 mM EDTA in PBS) and resuspended in FACS buffer containing the viability marker 7-AAD (1:100 BD Pharmingen) before filtering through a 70-µm strainer.

For the intracellular measurement of IL-1α and IL-1β, single-cell suspensions were pre-cultured in DMEM with 2.5% FBS, 100 µg/mL Penicillin and Streptomycin and stimulated with BD GolgiStop™ (1:1000, BD Biosciences) for 4 h in 5% $CO_2$ at 37°C followed by a pre-incubation with the viability dye eFluor 506 (1:400 eBioscience) for 20 min at 4°C. Then cell suspensions were washed, blocked with FcR Blocking Reagent (1:100 BD Pharmingen) and stained with surface antibodies (see Methods Table 1 for antibodies list) as described above. After a washing step with FACS buffer, cells were incubated for 45 min in Fix/Perm buffer (eBioscience, Invitrogen, ThermoFisher Scientific), followed by 5 min incubation in 1X Permeabilization buffer (eBioscience, Invitrogen, ThermoFisher Scientific). Next, the cells were stained for a minimum of 1 h in 1X Permeabilization buffer containing FcR Blocking Reagent (1:600 BD Pharmingen) and intracellular markers (see Methods Table 1 for antibody details). Cells were subsequently washed and resuspended in Permeabilization buffer before filtering through a 70-µm strainer.

For cell counting goals, counting beads (1:100 Spherotech) were added per sample. Flow cytometry analyses were performed on a BD Symphony A5 Cell Analyzer (BD Biosciences) and subsequently analysed using FlowJo v.10.6.1.

**Methods Table 1.** FACS Antibodies

| Anti | Conjugate | Company | Cat. No. | Clone |
|---|---|---|---|---|
| ARG1 | Pe-Cy7 | eBioscience | 25-3697-82 | A1efF5 |
| C1Q | FITC | Tebubio | 7501F | RmC7H8 |
| CD11b | BUV395 | BD Horizon | 563553 | M1/70 |
| CD11b | PE-Cy7 | BD Pharmingen | 552850 | M1/70 |
| CD19 | PE-Cy5 | eBioscience | 15-0193-82 | eBio1D3 |
| CD3 | Alexa Fluor 700 | BioLegend | 100215 | 17A2 |
| CD31 | BV421 | BioLegend | 102423 | 390 |
| CD326 | PE-Cy7 | BioLegend | 118215 | G8.8 |
| CD3e | eFluor 450 | eBioscience | 48-0032-82 | 17A2 |
| CD4 | BV605 | BioLegend | 100548 | RM4-5 |
| CD44 | AF700 | BioLegend | 103026 | IM7 |
| CD45 | BUV805 | BD OptiBuild | 748370 | 30-F11 |
| CD45 | APC-eFluor 780 | eBioscience | 47-0451-82 | 30-F11 |
| CD64 | BV711 | BioLegend | 139311 | X54-5/7.1 |
| CD8a | APC-Cy7 | eBioscience | 25-5773-82 | FJK-16s |
| FOXP3 | PE-Cy7 | eBioscience | 12-5773-82 | FJK16S |
| GP38 (PDPN) | Alexa Fluor 488 | BioLegend | 127405 | 8.1.1 |
| IL-1α | PE | BioLegend | 503203 | ALF-161 |
| IL-1β (Pro-form) | APC | eBioscience | 17-7114-80 | NJTEN3 |
| Live Dead | eFluor 506 | eBioscience | 65-0866-14 | |
| Live Dead | 7-AAD | BD Pharmingen | 51-68981E | |
| Ly6C | BV421 | BioLegend | 128043 | HK1.4 |
| Ly6C | BV650 | BioLegend | 128049 | HK1.4 |
| Ly6C | FITC | BD Pharmingen | 553104 | AL-21 |
| Ly6G | BUV563 | BD Horizon | 612921 | IA8 |
| Ly6G | APC | BD Pharmingen | 560599 | 1A8 |
| MHCII | APC-eFluor 780 | eBioscience | 47-5321-82 | M5/114.15.2 |
| MHCII | BV510 | BioLegend | 107636 | M5/114.15.2 |
| SiglecF | eFluor 660 | eBioscience | 50-1702-80 | 1RNM44N |

| | | | |
|---|---|---|---|
| SPP1 | PE | R&D systems | IC808P |

**Tumor-infiltrating monocyte sorting for EGCs stimulation.** Tumor-infiltrating monocytes were sorted from orthotopic tumors based on the expression of the viability marker 7-AAD, CD45, CD64, Ly6C, MHCII, SiglecF and Ly6G (see Methods Table 1 for antibodies details) using a Sony MA9000 sorter. Next, $1x10^5$ tumor or BM-derived monocytes were cultured in complete DMEM medium overnight in 5% $CO_2$ at 37°C. The conditioned medium of these monocytes was collected and used to stimulate primary embryonic neurosphere-derived EGCs ($5x10^4$ cells/mL) in the presence of 5 µg/mL IgG (BioXCell) or 5 µg/mL anti-IL-1R (BioXCell) for 24h in 5% $CO_2$ at 37°C.

**RNA extraction and gene expression**. RNA was isolated using the innuPREP RNA Mini Kit (Analytik Jena) or RNeasy Mini Kit (Qiagen) for tissue and high cell numbers or RNeasy Plus Micro Kit (Qiagen) for low cell numbers according to the manufacturer's instructions. Dependent on RNA concentrations, total RNA was transcribed into cDNA by the qScript™ cDNA SuperMix (QiantaBio) or the High-Capacity cDNA Reverse Transcription Kit (ThermoFisher Scientific) according to manufacturer's instructions. RT-PCR was performed with the LightCycler 480 SYBR Green I Master (Roche) on the Light Cycler 480 (Roche). Results were quantified using the $2^{-\Delta Ct}$ method. The expression levels of the genes of interest were normalized to the expression levels of the reference gene *Rpl32*.

**Methods Table 2**. Primer Sequences

| | Forward primer | Reverse primer |
|---|---|---|
| *Arg1* | CAGAAGAATGGAAGAGTCAG | CAGATATGCAGGGAGTCACC |
| *C1qa* | ATCCAGTTTGATCGGACCAC | CATCTTCAGCCACTGTCCATA |
| *Ccl2* | CAGGTGTCCCAAAGAAGCTGTA | CATTTGGTTCCGATCCAGG |
| *Ifng* | TCAAGTGGCATAGATGTGGAAGAA | TGGCTCTGCAGGATTTTCATG |
| *Il1a* | GAGAGCCGGGTGACAGTATC | ACTTCTGCCTGACGAGCTTC |
| *Il1b* | TTGACGGACCCCAAAAGATG | AGGACAGCCCAGGTCAAAG |
| *Il6* | CCATAGCTACCTGGAGTACATG | TGGAAATTGGGGTAGGAAGGAC |
| *Lcn2* | CGGAGCGATCAGTTCCGGG | GCCCTGGTCCTGGTCCCTGA |
| *Rpl32* | AAGCGAAACTGGCGGAAAC | TAACCGATGTTGGGCATCAG |
| *Spp1* | AGCAAGAAACTCTTCCAAGCAA | GTGAGATTCGTCAGATTCATCCG |
| *Timp1* | AGACAGCCTTCTGCAACT | CAGCCTTGAATCCTTTTAGCATC |
| *Tnfa* | CTATGGCCCAGACCCTCACACTC | GCTGGCACCACTAGTTGGTTGTCTT |

**Bulk RNA sequencing.** For Bulk RNA-seq of the *in vitro* tumor EGCs model, total RNA from *in vitro* generated unstimulated, H-CM and TME-CM primary embryonic neurosphere-derived EGCs was provided to the Genomics core (KU Leuven). QuantSeq 3' mRNA libraryprep (015, Lexogen) was used to generate cDNA libraries, followed by sequencing on the HiSeq4000 system. Quality control of raw reads was performed with FastQC v0.11.7 (Andrews S. FastQC: a quality control tool for high throughput sequence data. Available online at: http://www.bioinformatics.babraham.ac.uk/projects/fastqc, 2010.). Adapters were filtered with ea-utils fastq-mcf v1.05 (Erik Aronesty. ea-utils: Command-line tools for processing biological sequencing data. Available online at: https://github.com/ExpressionAnalysis/ea-utils, 2011.). Splice-aware alignment was performed with HISAT2 (Kim et al., 2019) against the reference genome mm10 using the default parameters. Reads mapping to multiple loci in the reference genome were discarded. Resulting BAM alignment files were handled with Samtools v1.5 (Li et al., 2009). Quantification of reads per gene was performed with HT-seq Count v0.10.0, Python v2.7.14 (Anders et al., 2015). Count-based differential expression analysis was done with R-based (The R Foundation for Statistical Computing, Vienna, Austria) Bioconductor package DESeq2 (Love et al., 2014). Reported p-values were adjusted for multiple testing with the Benjamini-Hochberg procedure, which controls false discovery rate (FDR). Data visualization was prepared using ggplot2 R package (v3.4.1) or pheatmap (v1.0.12).

For 3'mRNA sequencing of naive and AOM/DSS-treated mice, isolated RNA was provided to the Genomics Core Facility of the University Hospital Bonn, which performed library preparation using QuantSeq FWD 3'mRNA-Seq kit (Lexogen) according to the manufacturer's instructions. Sequencing was performed on the NovaSeq6000 with a sequencing depth of 10M raw reads. Data were analyzed using PartekFlow software available from https://www.partek.com/partek-flow/#features. Visualisation was done with PartekFlow software and GraphPad Prism 6.

**Weighted gene correlation network analysis (WGCNA).** First, variance stabilizing transformation was performed on the bulk RNA-seq data generated from unstimulated, H-CM and TME-CM primary neurosphere-derived EGCs using the DESeq2 (Love et al., 2014) package in R (v4.2.2). Next, WGCNA(Langfelder and Horvath, 2008) was performed using the R package WGCNA (v1.72.1). To distinguish the modules with different expression patterns, a soft threshold power of 12, which was the lowest power for the scale-free topology fit index on 0.85, was selected to produce a hierarchical clustering tree (dendrogram). The function "blockwiseModules" was used for automatic block-wise network construction and module identification. The number of modules was detected automatically by the algorithm, with the number of genes in a module limited to between 30 and 5000 genes. The co-expression networks were created based on the similarity of expression patterns of genes and the networks were established by merging genes with similar co-expression patterns into modules.

**Single cell RNA sequencing of orthotopic murine Tumors.** Cell suspensions of orthotopic murine tumors were processed with a 10x Chromium Next GEM Single Cell 5' kit and loaded on a 10x chromium controller to create Single Cell Gel beads in Emulsion (GEM). A cDNA library was created using a 10x 5' library kit and was then paired-end sequenced on an Illumina Novaseq device following 10x's guidelines (https://www.10xgenomics.com/support/single-cell-immune-profiling/documentation/steps/sequencing/sequencing-requirements-for-single-cell-v-d-j). Sample demultiplexing and data analysis was performed using 10x's Cellranger suite (https://support.10xgenomics.com/single-cell-vdj/software/pipelines/latest/using/vdj) using the standard parameters.

**Single-cell RNA sequencing Clustering and Dimensionality reduction.** The count matrices obtained after pre-processing with Cellranger were concatenated to obtain a combined raw count matrix which was then analyzed using the Seurat R package (v3.1.3). Cells with less than 300 or more than 6000 genes and cells with more than 15% mitochondrial genes were discarded from the analysis. Normalization and scaling were done with default variables with top variable genes identified using FindVariableFeatures function. After principal component analysis, 1$^{st}$ 39 principal components were used based on the elbow plot for creating a nearest neighbor graph using FindNeighbours function in Seurat. After clustering at a resolution of 1, clusters were classified into immune and non-immune clusters. 6 small doublet clusters with markers of two or more distinct cell types were removed. Also, two clusters with low nUMI and lacking distinguishing markers of any cell types were also removed. A subset of Seurat Object with immune clusters alone was created and the same pipeline was followed from Normalization to Clustering (number of principal components used = 32). After clustering at resolution 1, the clusters were manually annotated inspired by Zhang et al. (Zhang et al., 2020). Clusters annotated as monocytes or macrophages were re-clustered similarly to identify the subclusters. These sub-clusters were annotated manually based on the expression of Ly6c2, Ccr2, H2-Ab1, Spp1, C1qa, Cx3cr1, and Mki67. Further to learn potential differentiation trajectory, Monocle-3 was used. (Parameters: n center = 300, minimal branch length = 10, nn.k = 20). Genes upregulated or downregulated in SPP1$^{+}$ TAMs compared to C1Q$^{+}$ TAMs were functionally annotated using universal enrichment function ‘enricher’ from the ‘ClusterProfiler’ package (v4.6.0) with a gene annotation database aggregation containing all terms from Human Phenotype, Transcription factor, and Hallmark from Molecular Signature Database (MSigDB), BIOCARTA, REACTOME, GO and KEGG Markers for different clusters were determined using a Wilcoxon rank sum test with FindMarkers or FindAllMarkers functions in Seurat.

**Inferring cell-cell communication using NicheNet.** NicheNet (nichenetr R package; v1.1.0) was used to study the interactions between EGCs and tumor-infiltrating monocytes. To identify TME EGC-derived ligands potentially inducing the differentiation of monocytes into SPP1$^{+}$ TAMs, bulk RNA-seq data from *in vitro* TME-CM EGCs was used. Ligands were identified after filtering for genes upregulated in 24h time point TME-CM EGCs with respect to 24h timepoint H-CM EGCs (adjusted p value < 0.05). Genes differentially expressed between SPP1⁺ TAMs and monocytes (adjusted p value < 0.05) were considered as the gene set of interest.

NicheNet was also used to study the interaction between tumor-infiltrating immune cells and EGCs. To identify immune cell-derived ligands potentially inducing differentiation of 24h time point H-CM EGCs into 24h time point TME-CM EGCs, scRNA-seq data of the immune compartment of the *in vivo* murine orthotopic CRC model was used. Using get_expressed_genes function from NichenetR, genes expressed in at least 5 % of immune cell clusters were considered as potential ligands. Genes differentially expressed between TME-CM EGCs and H-CM EGCs (adjusted p value < 0.001) were considered as the gene set of interest.

**Bio-informatic analysis**: **KUL3 Dataset.** Bio-informatic analysis of the CRC tumor microenvironment of patients affected by CRC was performed making use of the published KUL3 dataset (Lee et al., 2020). Integration of the data, dimensionality reduction, unsupervised clustering and differential gene expression analysis was performed in R using Seurat with SCTransform - Integration pipeline. For downstream analysis, "border" and "tumor" samples were taken together and considered as tumor samples. Patient KUL31 was excluded from all EGCs analysis, due to extremely low cell numbers. Pathway enrichment analysis was done using Ingenuity pathway Analysis (IPA, Qiagen). Modules identified using WGCNA on mouse bulk RNA-seq data was converted to one-to-one human orthologs and then used for single-sample Gene Set Enrichment Analysis GSEA (ssGSEA) as implemented in single-cell Gene Set Variation Analysis (scGSVA) R package (v0.0.11).

**TCGA analysis: data acquisition, patients' classification and survival analysis.** The processed gene expression RNA-seq (IlluminaHiSeq) data of the Cancer Genome Atlas (TCGA) colorectal adenocarcinoma (COADREAD) was downloaded from University of California SantaCruz (UCSC) Xena using the UCSCXenaTools (ref: https://joss.theoj.org/papers/10.21105/joss.01627) R library. The details of data integration and processing are described in UCSC-Xena browser (https://xenabrowser.net/). The clinical information and overall survival (OS) data of the patients were also obtained using UCSCXenaTools (data subtype: "phenotype"). According to their age, the patients were classified as above 65 years (>=65) and below 65 years. Patients with tumor stage I and IA were clustered as stage I, patients with stage II, IIA, IIB as stage II, patients with stage III, IIIA, IIIB, IIIC as stage III, and patients with stage IV, IVA, IVB as stage IV. Patients with microsatellite stability were classified as MSS and patients with microsatellite instability high and low as MSI. The consensus molecular subtypes (CMS) were predicted using the R package CMSClassifier (v1.0.0) and the intrinsic CMS (iCMS) classification of the patients was performed as previously described (Joanito et al., 2022). The 376 patients affected by CRC were hierarchically clustered according to the high and low expression patterns of the specific gene signatures (Methods Table 3) in the tumor samples. The R packages survival (v3.5.5) and survminer (v0.4.9) were used for survival analysis and plotting the Kaplan-Meier (KM) survival curve. A statistically significant difference in survival was indicated by a log-rank test p-value of $p<0.05$. Survival analysis with univariate and multivariate proportional hazards regression models (Cox regression) was performed to adjust for age, gender, radiation therapy, stage and EGC signature expression. The R packages pheatmap, gtsummary, and ggplot2 were used for visualization.

**Methods Table 3**

| | Gene signature |
|---|---|
| Enteric glial cells (manually curated from literature (Lee et al., 2020; Drokhlyansky et al., 2020; Kinchen et al., 2018)) | S100B, SOX10, PLP1, GFAP, CRYAB, CLU, FXYD1, ALDH1A1, PMP22, CDH19, SCN7A, PRNP |
| $SPP1^{+}$ TAMs (Zhang et al., 2020) | SPP1, PCSK5, SLC11A1, VCAN, SLC25A37, FLNA, UPP1, BCL6, AQP9, TIMP1, VEGFA, ADM, MARCO, FN1, IL1RN |
| Gliosis (Schneider et al., 2022) | PTGS2, CD44, CSF1, GCNT2, TGFB1, TUBB6, ICAM1, TNC, TNFRSF12A, RFC3, CP, HMOX1, EPHA4, PVR, PTX3, VCAN, CCL2, CTSB, THBS1, SRXN1, CXCL1, SPHK1, TIMP1, HMGA1, IL6, CXCL2, GFAP, NES, GDNF |

**Data availability.** ScRNA-seq and bulk RNA-seq data generated for this study will be deposited in the Gene Expression Omnibus (GEO) database and will be made publicly available.

**Code availability.** No custom algorithms were used in the analysis. Code for any specific analysis is available from authors upon request.

Pombo Antunes, A.R., I. Scheyltjens, F. Lodi, J. Messiaen, A. Antoranz, J. Duerinck, D. Kancheva, L. Martens, K. De Vlaminck, H. Van Hove, S.S. Kjølner Hansen, F.M. Bosisio, K. Van der Borght, S. De Vleeschouwer, R. Sciot, L. Bouwens, M. Verfaillie, N. Vandamme, R.E. Vandenbroucke, O. De Wever, Y. Saeys, M. Guilliams, C. Gysemans, B. Neyns, F. De Smet, D. Lambrechts, J.A. Van Ginderachter, and K. Movahedi. 2021. Single-cell profiling of myeloid cells in glioblastoma across species and disease stage reveals macrophage competition and specialization. *Nat. Neurosci.* 24:595–610. doi:10.1038/s41593-020-00789-y.

Ritchie, M.E., B. Phipson, D. Wu, Y. Hu, C.W. Law, W. Shi, and G.K. Smyth. 2015. Limma powers differential expression analyses for RNA-sequencing and microarray studies. *Nucleic Acids Res.* 43:e47. doi:10.1093/nar/gkv007.

Schneider, R., P. Leven, S. Mallesh, M. Breßer, L. Schneider, E. Mazzotta, P. Fadda, T. Glowka, T.O. Vilz, P. Lingohr, J.C. Kalff, F.L. Christofi, and S. Wehner. 2022. IL-1-dependent enteric gliosis guides intestinal inflammation and dysmotility and modulates macrophage function. *Commun. Biol.* 5:1–16. doi:10.1038/s42003-022-03772-4.

Stakenborg, M., S. Abdurahiman, V. De Simone, G. Goverse, N. Stakenborg, L. Van Baarle, Q. Wu, D. Pirottin, J. Kim, L. Chappell-maor, I. Pintelon, E. Pollenus, L. Boon, P. Van Den Steen, M. Hao, J.A. Van Ginderachter, G.E. Boeckxstaens, J. Timmermans, and S. Jung. 2022. Enteric glial cells favor accumulation of anti-inflammatory macrophages during the resolution of muscularis in flammation. *Mucosal Immunol.* 15:1296–1308. doi:10.1038/s41385-022-00563-2.

The, M., M.J. MacCoss, W.S. Noble, and L. Käll. 2016. Fast and Accurate Protein False Discovery Rates on Large-Scale Proteomics Data Sets with Percolator 3.0. *J. Am. Soc. Mass Spectrom.* 27:1719–1727. doi:10.1007/s13361-016-1460-7.

Zhang, L., Z. Li, K.M. Skrzypczynska, Q. Fang, W. Zhang, S.A. O’Brien, Y. He, L. Wang, Q. Zhang, A. Kim, R. Gao, J. Orf, T. Wang, D. Sawant, J. Kang, D. Bhatt, D. Lu, C.M. Li, A.S. Rapaport, K. Perez, Y. Ye, S. Wang, X. Hu, X. Ren, W. Ouyang, Z. Shen, J.G. Egen, Z. Zhang, and X. Yu. 2020. Single-Cell Analyses Inform Mechanisms of Myeloid-Targeted Therapies in Colon Cancer. *Cell*. 181:442–459. doi:10.1016/j.cell.2020.03.048.

**SUPPLEMENTAL FIGURES TITLES AND LEGENDS**

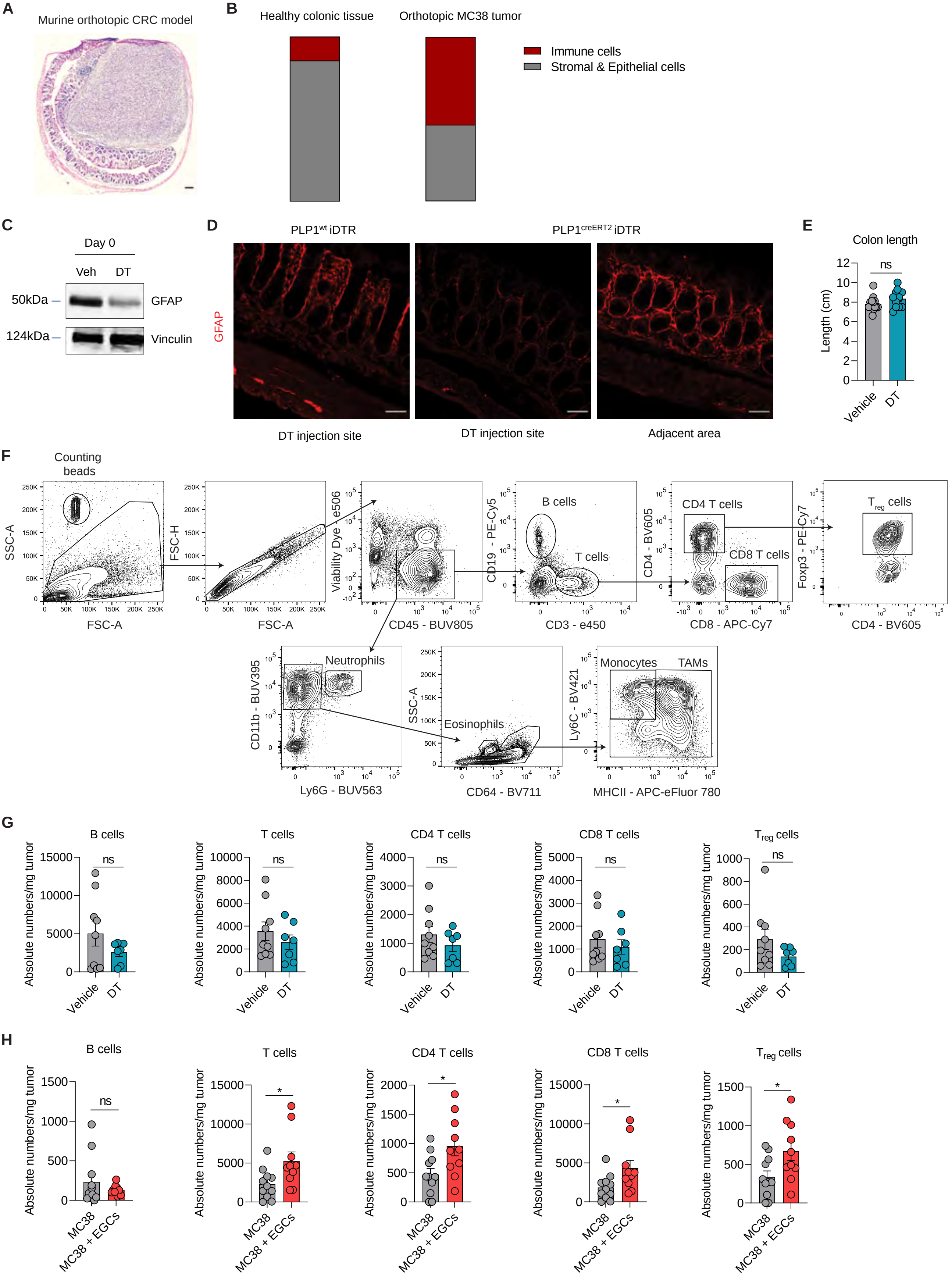

A
Murine orthotopic CRC model
B
Healthy colonic tissue
Orthotopic MC38 tumor
Immune cells
Stromal & Epithelial cells
C
Day 0
Veh
DT
50kDa
GFAP
124kDa
Vinculin
D
PLP1wt iDTR
PLP1creERT2 iDTR
GFAP
DT injection site
DT injection site
Adjacent area
E
Colon length
Length (cm)
ns
Vehicle
DT
F
Counting beads
SSC-A
FSC-A
FSC-H
Viability Dye - e506
CD45 - BUV805
CD19 - PE-Cy5
B cells
T cells
CD3 - e450
CD4 - BV605
CD4 T cells
CD8 T cells
CD8 - APC-Cy7
Foxp3 - PE-Cy7
Treg cells
CD11b - BUV395
Neutrophils
Ly6G - BUV563
Eosinophils
CD64 - BV711
Ly6C - BV421
Monocytes
TAMs
MHCII - APC-eFluor 780
G
B cells
T cells
CD4 T cells
CD8 T cells
Treg cells
Absolute numbers/mg tumor
ns
Vehicle
DT
H
B cells
T cells
CD4 T cells
CD8 T cells
Treg cells
Absolute numbers/mg tumor
ns
*
MC38
MC38 + EGCs

**Figure S1. EGCs dictate the immune populations in the TME**

(A-B) BL6 mice were injected endoscopically in the colonic submucosa at day(d)0 with MC38 cells. Tumor growth and cellular infiltration were assessed at d21. representative hematoxylin and eosin image of orthotopic murine tumor section, scale bar 500 µm (A). Abundance presentation of immune or stromal and epithelial cells in healthy and orthotopic CRC tissues (B).

(C) Representative western blots for GFAP and Vinculin at d0 of total colonic tissue from $PLP1^{CreERT2}$iDTR mice intracolonic (i.c.) injected at d-5 and d-3 with 40 ng Diphtheria toxin (DT) or saline (Veh).

(D) Representative image showing GFAP (red) staining in the colon at d0 of $PLP1^{Wt}$iDTR and $PLP1^{CreERT2}$iDTR mice i.c. injected at d-5 and d-3 with 40 ng Diphtheria toxin (DT), scale bar 50 µm.

(E-G) $PLP1^{CreERT2}$iDTR mice were i.c. injected at d-5 and d-3 with 40 ng Diphtheria toxin (DT) or saline (Vehicle) ($n = 9$ Vehicle, $n = 7$ DT). On d0 both groups were i.c. injected with MC38 cells and on d7 the colon length (E) and tumor-infiltrating lymphoid immune cells were assessed. FACS gating strategy to identify the lymphoid (up) and myeloid (down) populations in the tumor microenvironment (F). Data of immune cells are presented as absolute numbers per mg of tumor tissue (G).

(H) BL6 mice were i.c. injected at d0 with MC38 cells and with or without embryonic neurosphere-derived EGCs (1:1 ratio). The tumor-infiltrating lymphoid immune cells were assessed by FACS on d21. Data are presented as absolute numbers per mg of tumor tissue ($n = 11$ MC38, $n = 10$ MC38 + EGCs).

Data are represented as mean ± SEM. Statistical analysis: unpaired Mann-Whitney (E, G-H) *$p < 0.05$, ns not significant.

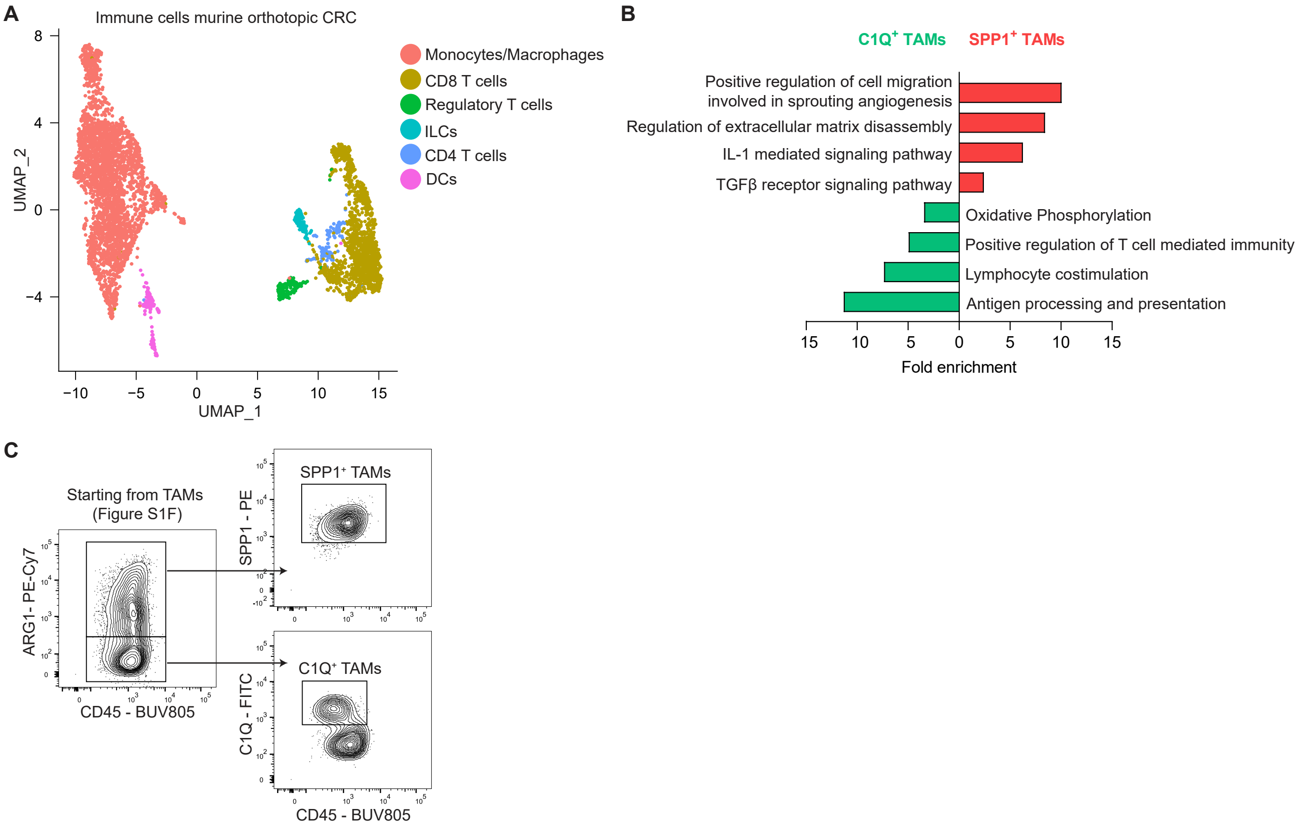
A
Immune cells murine orthotopic CRC
Monocytes/Macrophages
CD8 T cells
Regulatory T cells
ILCs
CD4 T cells
DCs
UMAP_2
UMAP_1
B
C1Q+ TAMs
SPP1+ TAMs
Positive regulation of cell migration involved in sprouting angiogenesis
Regulation of extracellular matrix disassembly
IL-1 mediated signaling pathway
TGFβ receptor signaling pathway
Oxidative Phosphorylation
Positive regulation of T cell mediated immunity
Lymphocyte costimulation
Antigen processing and presentation
Fold enrichment
C
Starting from TAMs (Figure S1F)
ARG1- PE-Cy7
CD45 - BUV805
SPP1+ TAMs
SPP1 - PE
C1Q+ TAMs
C1Q - FITC
CD45 - BUV805

**Figure S2. Dichotomy of functional phenotypes of TAMs in murine orthotopic CRC**

(A) UMAP of scRNA-seq data from tumor-infiltrating immune cells in BL6 mice bearing orthotopic colon tumors, d21 after tumor induction *(n = 3).*

(B) Gene ontology biological process (GOBP) analysis showing the differential pathways enriched in $SPP1^+$ TAMs versus $C1Q^+$ TAMs, data extracted from scRNAseq murine orthotopic CRC dataset.

(C) $SPP1^+$ and $C1Q^+$ TAMs FACS gating strategy.

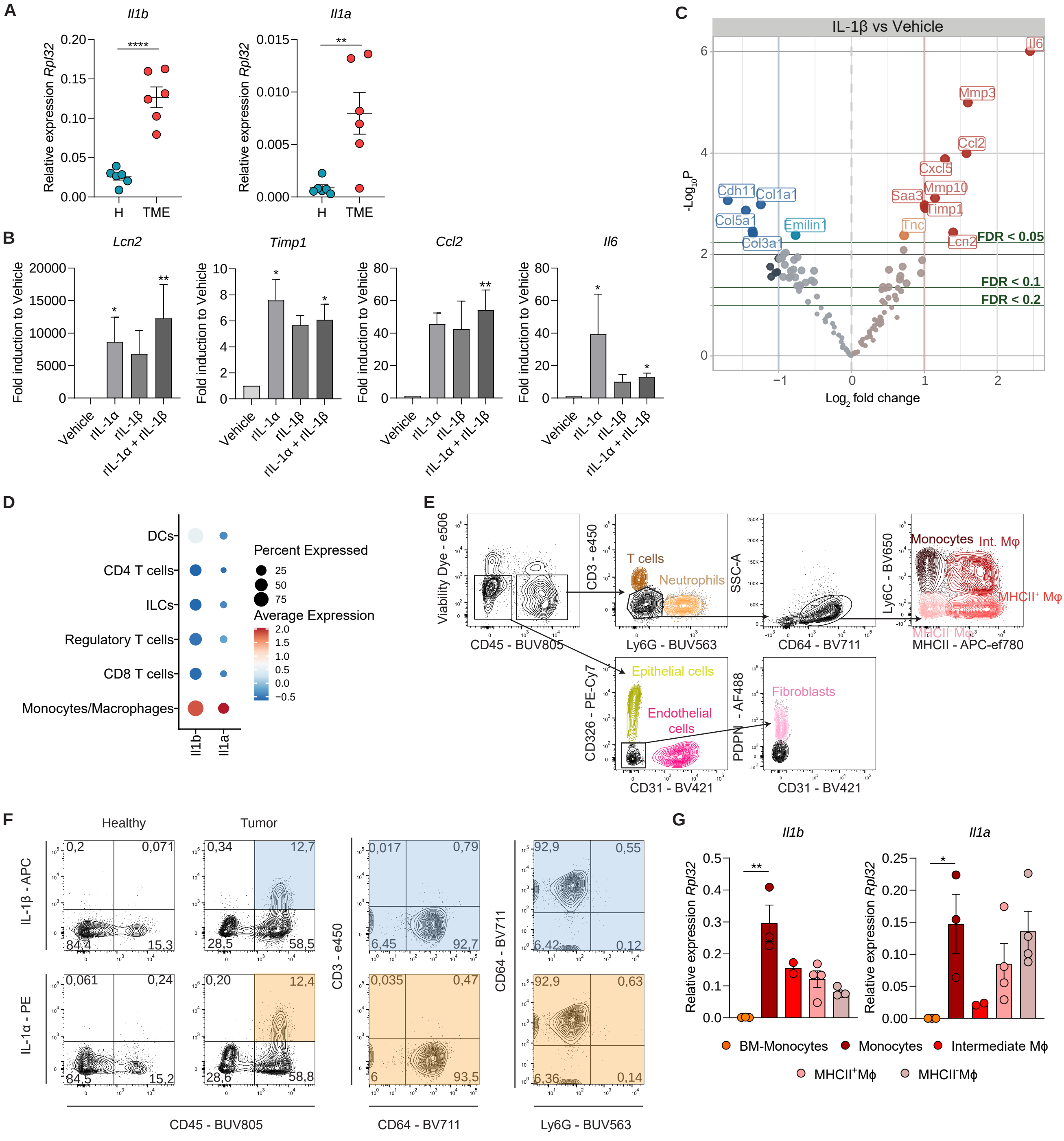

A
Il1b
Il1a
Relative expression Rpl32
H
TME
B
Lcn2
Timp1
Ccl2
Il6
Fold induction to Vehicle
Vehicle
rIL-1α
rIL-1β
rIL-1α + rIL-1β
C
IL-1β vs Vehicle
Il6
Mmp3
Ccl2
Cxcl5
Mmp10
Saa3
Timp1
Tnc
Lcn2
Cdh11
Col1a1
Col5a1
Emilin1
Col3a1
FDR < 0.05
FDR < 0.1
FDR < 0.2
-Log10P
Log2 fold change
D
DCs
CD4 T cells
ILCs
Regulatory T cells
CD8 T cells
Monocytes/Macrophages
Il1b
Il1a
Percent Expressed
Average Expression
E
Viability Dye - e506
CD45 - BUV805
CD3 - e450
Ly6G - BUV563
T cells
Neutrophils
SSC-A
CD64 - BV711
Ly6C - BV650
MHCII - APC-ef780
Monocytes
Int. Mφ
MHCII+ Mφ
MHCII- Mφ
CD326 - PE-Cy7
CD31 - BV421
Epithelial cells
Endothelial cells
PDPN - AF488
Fibroblasts
F
Healthy
Tumor
IL-1β - APC
IL-1α - PE
CD3 - e450
CD64 - BV711
CD45 - BUV805
CD64 - BV711
Ly6G - BUV563
G
Il1b
Il1a
Relative expression Rpl32
BM-Monocytes
Monocytes
Intermediate Mφ
MHCII+Mφ
MHCII-Mφ

**Figure S3. TME-derived IL-1 induces the CRC EGCs signature *in vitro***

(A) Relative mRNA levels for *Il1b* and *Il1a* normalized to the housekeeping gene *Rpl32,* in a single-cell suspension of digested murine healthy colon (H) or orthotopic tumor microenvironment (TME) tissue *(n = 6)*.

(B) Relative mRNA levels for *Lcn2, Timp1, Ccl2*, and *Il6* in primary embryonic neurosphere-derived EGCs stimulated for 24h with or without recombinant (r) IL-1α (10 ng/mL), IL-1β (10 ng/mL) or the combination of the two (*n = 6* vehicle, *n = 3* rIL-1α and rIL-1β, *n = 6* rIL-1α + rIL-1β).

(C) Volcano plot of differentially expressed proteins between primary adult neurosphere-derived EGCs treated for 24h with rIL-1β (10 ng/ml) or Vehicle. Protein concentration in the culture supernatants was determined by liquid chromatography/mass spectrometry (*n = 4*).

(D) Dot plot showing expression of *Il1b* and *Il1a* in the tumor-infiltrating immune cell clusters identified by scRNA-seq analysis of mice bearing orthotopic colon tumors *(n = 3)*.

(E) FACS gating strategy to identify immune and stromal populations in the TME (Int. MΦ, intermediate Macrophages).

(F) Contour plots representing IL-1β and IL-1α expression in healthy colon and orthotopic CRC tumors based on CD45 expression (left), CD3 and CD64 expression (middle) and CD64 and Ly6G expression (right).

(G) Relative mRNA levels for *Il1b* and *Il1a* normalized to the housekeeping gene *Rpl32* in tumor-infiltrating monocytes, intermediate macrophages (MΦ), $MHCII^{+}$ MΦ and $MHCII^{-}$ MΦ and bone marrow (BM)-derived monocytes from mice bearing orthotopic colon tumors (*n = 3* BM-derived monocytes and tumor-infiltrating monocytes, *n = 2* intermediate MΦ and *n = 4* $MHCII^{+}$ MΦ and $MHCII^{-}$ MΦ).

All data are represented as mean ± SEM (A-B and G). Statistical analysis: unpaired t-test (A), Kruskal-Wallis test with correction for multiple comparisons, compared to Vehicle (B) or one-way ANOVA with correction for multiple comparisons (G). *p < 0.05, ** p <0.005, **** p <0.00005, ns not significant.

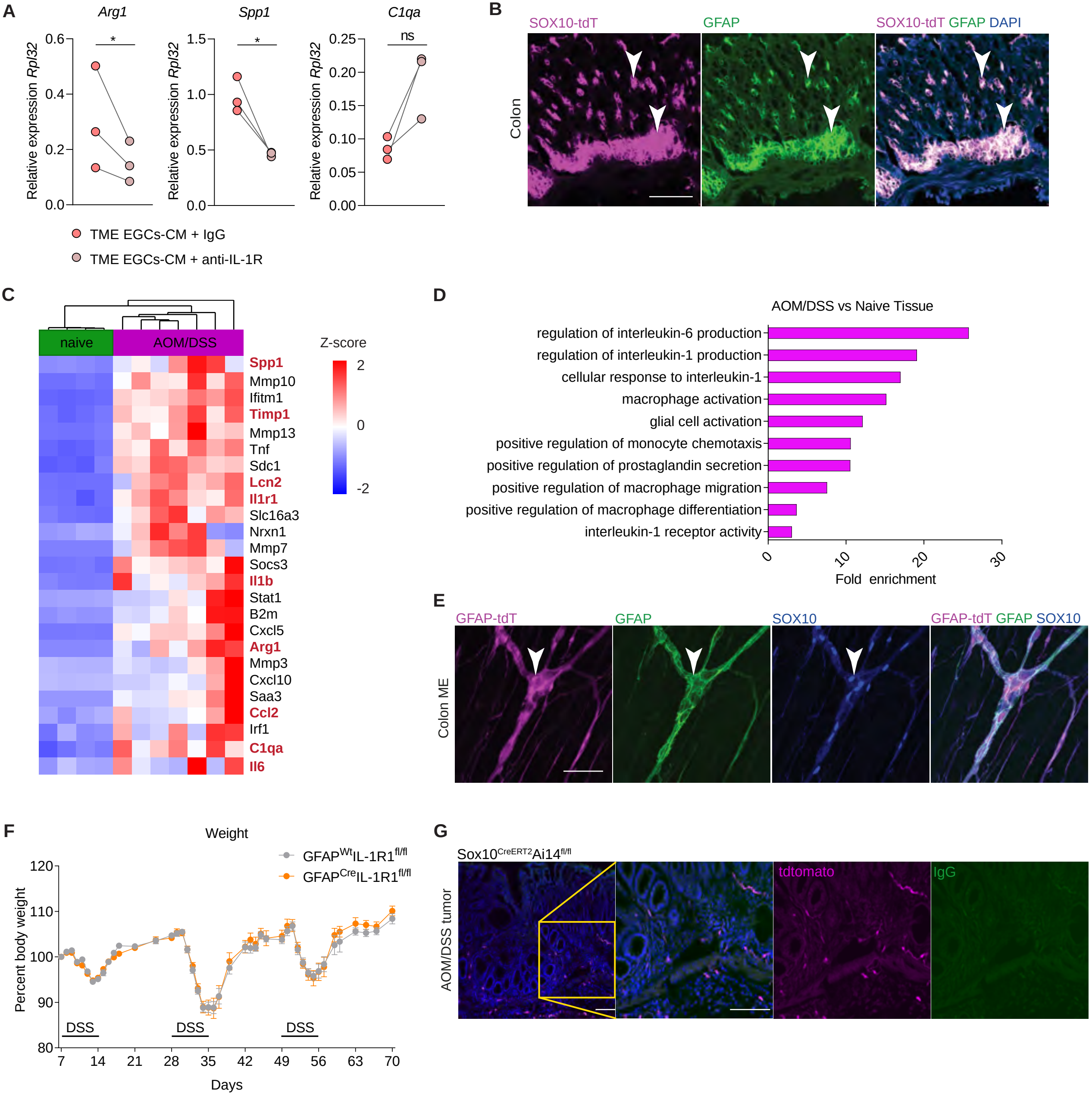
A
Arg1
Spp1
C1qa
Relative expression Rpl32
*
*
ns
0.6
0.4
0.2
0.0
1.5
1.0
0.5
0.0
0.25
0.20
0.15
0.10
0.05
0.00
TME EGCs-CM + IgG
TME EGCs-CM + anti-IL-1R
B
SOX10-tdT
GFAP
SOX10-tdT GFAP DAPI
Colon
C
naive
AOM/DSS
Z-score
2
0
-2
Spp1
Mmp10
Ifitm1
Timp1
Mmp13
Tnf
Sdc1
Lcn2
Il1r1
Slc16a3
Nrxn1
Mmp7
Socs3
Il1b
Stat1
B2m
Cxcl5
Arg1
Mmp3
Cxcl10
Saa3
Ccl2
Irf1
C1qa
Il6
D
AOM/DSS vs Naive Tissue
regulation of interleukin-6 production
regulation of interleukin-1 production
cellular response to interleukin-1
macrophage activation
glial cell activation
positive regulation of monocyte chemotaxis
positive regulation of prostaglandin secretion
positive regulation of macrophage migration
positive regulation of macrophage differentiation
interleukin-1 receptor activity
0
10
20
30
Fold enrichment
E
GFAP-tdT
GFAP
SOX10
GFAP-tdT GFAP SOX10
Colon ME
F
Weight
GFAP^Wt IL-1R1^fl/fl
GFAP^Cre IL-1R1^fl/fl
Percent body weight
120
110
100
90
80
DSS
DSS
DSS
7
14
21
28
35
42
49
56
63
70
Days
G
Sox10^CreERT2 Ai14^fl/fl
AOM/DSS tumor
tdtomato
IgG

**Figure S4. IL-1R activation in EGCs promotes SPP1$^+$ TAM differentiation *in vitro* and *in vivo***

(A) Relative mRNA levels for *Arg1, Spp1* and *C1qa,* normalized to the housekeeping gene *Rpl32,* in murine bone marrow-derived monocytes cultured for 48h with supernatant of primary embryonic neurosphere-derived EGCs, which were pre-incubated for 24h with tumor microenvironment conditioned medium (TME-CM) together with isotype IgG (5 µg/mL) or anti-IL-1R (5 µg/mL). *(n = 3).*

(B) Representative image of the EGC reporter line Sox10$^{CreERT2}$Ai14$^{fl/fl}$ showing tdtomato (magenta), GFAP (green), and DAPI (blue) in colon tissue section (scale bar 50 µm).

(C-D) Wildtype mice were subjected to the AOM/DSS model as described in Figure 6D. Naive and tumor tissues were collected at d70 and processed for 3'bulk mRNA-sequencing *(n = 7* AOM/DSS, *n = 4* naive). Heatmap of selected genes upregulated in AOM/DSS compared to naive samples (C). Gene set enrichment analysis for the differentially upregulated genes in AOM/DSS-treated compared to naive mice (D).

(E) Representative image of the EGC reporter line GFAP$^{Cre}$Ai14$^{fl/fl}$ showing tdtomato (magenta), GFAP (green), and SOX10 (blue) in a colon muscularis whole mount (scale bar 50 µm).

(F) GFAP$^{Wt}$IL-1R1$^{fl/fl}$ and GFAP$^{Cre}$IL-1R1$^{fl/fl}$ littermates were subjected to the AOM/DSS model as described in Figure 6D. Weight curve of AOM/DSS-treated GFAP$^{Wt}$IL-1R1$^{fl/fl}$ (*n = 29* d7- 43, *n = 18* d44-70) and GFAP$^{Cre}$IL-1R1$^{fl/fl}$ mice (*n = 25* d7- 43, *n = 16* d44-70)

(G) Sox10$^{CreERT2}$Ai14$^{fl/fl}$ mice were subjected to the AOM/DSS model as described in Figure 6D. Representative image of tdtomato (magenta), IgG (green), and DAPI (blue) in tumor section at d70 (scale bar 100 µm).

Statistical analysis: paired t-test (A). *p < 0.05, ns not significant.

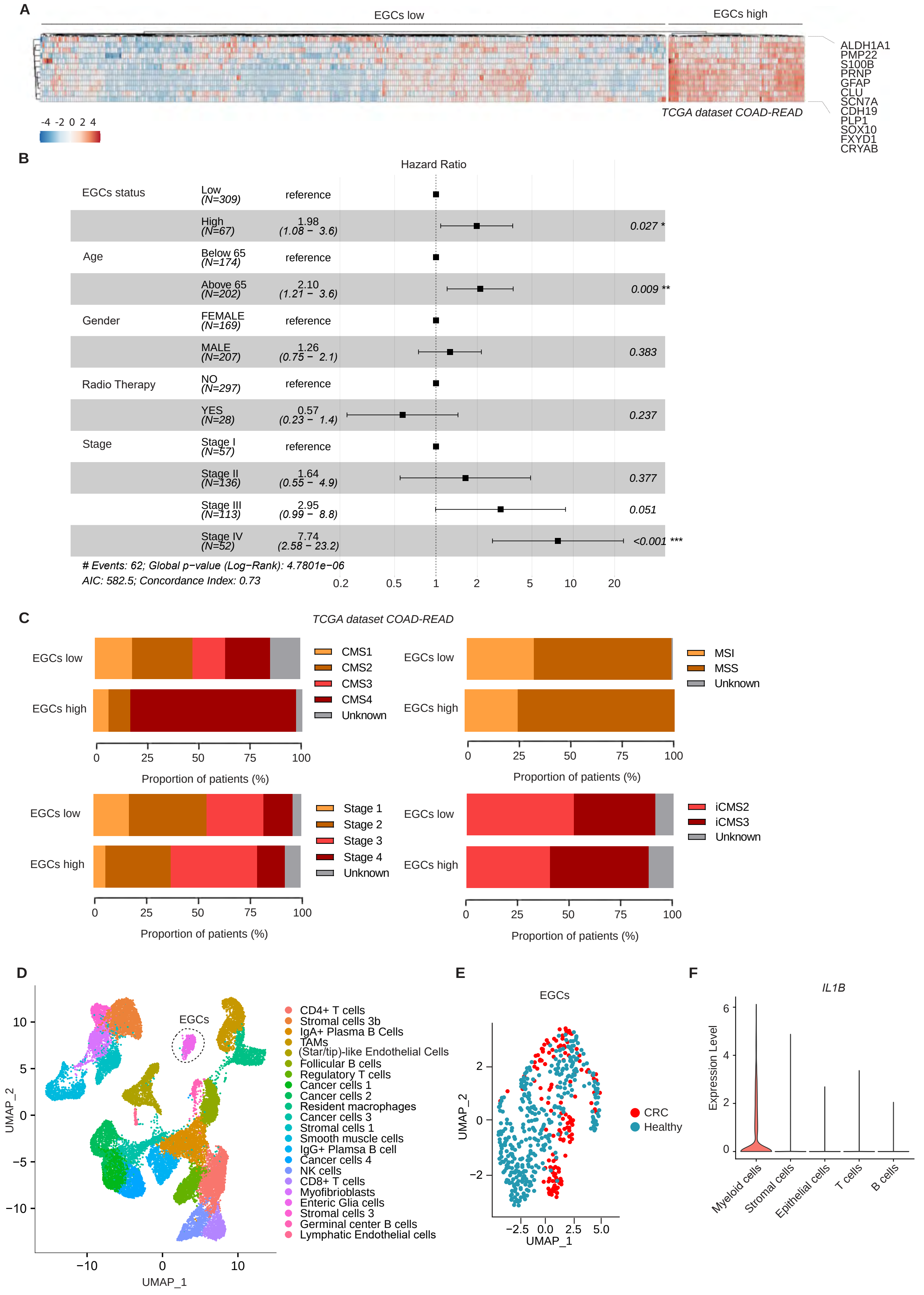
A
EGCs low
EGCs high
ALDH1A1
PMP22
S100B
PRNP
GFAP
CLU
SCN7A
CDH19
PLP1
SOX10
FXYD1
CRYAB
TCGA dataset COAD-READ
-4 -2 0 2 4
B
Hazard Ratio
EGCs status
Low (N=309) reference
High (N=67) 1.98 (1.08 − 3.6) 0.027 *
Age
Below 65 (N=174) reference
Above 65 (N=202) 2.10 (1.21 − 3.6) 0.009 **
Gender
FEMALE (N=169) reference
MALE (N=207) 1.26 (0.75 − 2.1) 0.383
Radio Therapy
NO (N=297) reference
YES (N=28) 0.57 (0.23 − 1.4) 0.237
Stage
Stage I (N=57) reference
Stage II (N=136) 1.64 (0.55 − 4.9) 0.377
Stage III (N=113) 2.95 (0.99 − 8.8) 0.051
Stage IV (N=52) 7.74 (2.58 − 23.2) <0.001 ***
### Events: 62; Global p−value (Log−Rank): 4.7801e−06
AIC: 582.5; Concordance Index: 0.73
0.2 0.5 1 2 5 10 20
C
TCGA dataset COAD-READ
EGCs low
EGCs high
CMS1
CMS2
CMS3
CMS4
Unknown
MSI
MSS
Unknown
Stage 1
Stage 2
Stage 3
Stage 4
Unknown
iCMS2
iCMS3
Unknown
0 25 50 75 100
Proportion of patients (%)
D
EGCs
UMAP_1
UMAP_2
CD4+ T cells
Stromal cells 3b
IgA+ Plasma B Cells
TAMs
(Star/tip)-like Endothelial Cells
Follicular B cells
Regulatory T cells
Cancer cells 1
Cancer cells 2
Resident macrophages
Cancer cells 3
Stromal cells 1
Smooth muscle cells
IgG+ Plamsa B cell
Cancer cells 4
NK cells
CD8+ T cells
Myofibroblasts
Enteric Glia cells
Stromal cells 3
Germinal center B cells
Lymphatic Endothelial cells
E
EGCs
CRC
Healthy
UMAP_1
UMAP_2
F
IL1B
Expression Level
Myeloid cells
Stromal cells
Epithelial cells
T cells
B cells

**Figure S5. EGCs are enriched in CRC tumors of patients with CMS4**

(A-C) TCGA COAD and READ patients stratified based on their expression of the EGCs signature genes (*n = 309* EGCs low, *n = 67* EGCs high). Heatmap of patients clustering (A). Cox logistic regression multivariate analysis of overall survival in CRC patients according to expression of EGCs signature genes and all the other relevant clinical parameters. For each variable, the reference level is the first one, P values indicate association with prognosis in this multivariate model. Error bars represent the 95% confidence interval (B). The proportion of the EGCs high and low patients classified in the different disease stages (top left), MSI/MSS (top right) CMS subtypes (bottom left) and iCMS subtypes (bottom right) (C).

(D-F) Transcriptome analysis of human CRC and healthy colon tissues in the KUL3 Dataset, Lee H. O. et al. 2020 (*n = 5*). UMAP of full scRNA-seq dataset, indicating the EGCs cluster (D). UMAP of isolated EGCs cluster (E). Violin plot showing expression of *IL1B* in the tumor micro-environment clusters (F).

**Table S1.** Differentially expressed genes and GO terms in Modules of weighted gene correlation network analysis (WGCNA), related to Figure 2C.

**Table S2.** Differentially expressed genes in EGCs transcriptomic analysis and significantly increased proteins in IL-1 stimulated EGCs mass spectrometry analysis. Related to Figures 2, 4 and 7.

**Table S3**. Differentially expressed genes between full-thickness AOM/DSS-treated colonic tumors and naive colonic tissues identified by 3'bulk mRNA-seq. Related to Figure S4.
